# Exercise-Induced Epoxy-Eicosanoids Promote Cardiomyocyte Proliferation and Post-Infarction Repair via NR4A1

**DOI:** 10.64898/2026.01.12.699155

**Authors:** Jin Wang, Zhongze Zhang, Yang Bian, Langran Wang, Mei Lan, Zhihua Zhang, Wenbin Cai, Xin Zhang, Yi Zhu, Xu Zhang

## Abstract

**BACKGROUND:** Myocardial infarction (MI) causes irreversible cardiomyocyte loss that exceeds the adult heart’s regenerative capacity, leading to adverse remodeling and heart failure. While exercise provides cardiovascular protection, the mechanisms underlying exercise-induced cardiac repair remain unclear.

**METHODS:** Plasma levels of free fatty acids and polyunsaturated fatty acid metabolites were quantified by liquid chromatography–tandem mass spectrometry (LC-MS/MS). RNA sequencing was performed to identify differentially expressed genes and enriched pathways. NR4A1 was knocked down in mice via tail vein injection of AAV9 carrying shRNA targeting NR4A1 (AAV9-shNR4A1). The role of 17,18-EEQ was evaluated by continuous subcutaneous infusion via osmotic minipump; the involvement of the cAMP–PKA–CREB axis was assessed using specific pharmacological inhibitor. Molecular analyses included ELISA, immunofluorescence staining, QPCR, and western blotting. Additionally, clinical samples were collected from 36 endurance-trained athletes and 30 sedentary individuals (age 18-22, male) for comparative analysis.

**RESULTS:** Here we demonstrate that preventive aerobic training promotes post-MI cardiomyocyte proliferation and confers cardioprotection by reducing fibrosis area and improving cardiac function. Targeted lipidomics revealed that exercise elevates circulating epoxyeicosatetraenoic acids (EEQs), particularly 17,18-EEQ; osmotic pump delivery of 17,18-EEQ mimicked the protective effects of exercise, reducing infarct burden and enhancing proliferation markers (Ki67, pH3, Aurora B). Transcriptomic analysis identified NR4A1 as the most prominent up-regulated mRNA under exercise induction; AAV9-mediated NR4A1 knockdown abolished exercise-mediated cardioprotection and proliferative benefits, establishing NR4A1 as essential for exercise-driven repair. Phosphoproteomic profiling revealed that 17,18-EEQ activates the PKA–CREB pathway to induce NR4A1 expression; pharmacological inhibition of adenylyl cyclase, PKA, or CREB phosphorylation reversed EEQ-mediated cardioprotection and proliferation, confirming pathway necessity. Dietary supplementation with eicosapentaenoic acid (EPA), the metabolic precursor of 17,18-EEQ, amplified exercise-induced cardioprotection by enhancing EEQ biosynthesis.

**CONCLUSION:** Our study identifies 17,18-EEQ as an exercise-derived exerkine that promotes cardiomyocyte proliferation and post-MI repair through a PKA–CREB–NR4A1 signaling axis, advancing our understanding of exercise cardioprotection and myocardial regeneration mechanisms while providing novel therapeutic targets for cardiac intervention.

**What IS New?:**

1. Aerobic exercise enhances post-myocardial infarction cardiac repair by upregulating the cardio-protective lipid mediator 17,18-EEQ and its downstream effector NR4A1.
2. 17,18-EEQ promotes cardiomyocyte proliferation and cardiac repair through activation of the cAMP–PKA–NR4A1 signaling axis.
3. Combining aerobic exercise with omega-3 fatty acid supplementation synergistically improves cardiac remodeling and functional recovery after ischemic injury.

**What Are the Clinical Implications?:**

1. Targeting endogenous cardiomyocyte proliferation represents a promising therapeutic strategy for cardiac regeneration after ischemic injury.
2. Dietary supplementation with omega-3 fatty acids may be a clinically feasible approach to amplify exercise-induced cardioprotection and improve outcomes after myocardial infarction.

## INTRODUCTION

Ischemic heart disease remains a leading global cause of death and disability, driven by cardiomyocyte loss, inflammation, and fibrosis that culminate in adverse remodeling and heart failure ^[1]^. Adult cardiomyocytes have limited proliferative potential—the annual renewal rate in humans is roughly 1% ^[2]^—yet recent studies demonstrate that modulating the cell cycle can promote structural and functional repair in models of myocardial infarction and ischemia–reperfusion, highlighting translational promise in cardiac regeneration ^[3]^. Metabolic reprogramming is a key determinant of postnatal cell-cycle exit; tuning glycolysis, fatty acid oxidation, and oxidative phosphorylation not only reshapes substrate utilization but also couples to cell-cycle programs through epigenetic pathways, thereby enhancing myocardial regeneration and improving cardiac function in adult injury models ^[4, 5]^. Notably, cardiomyocyte proliferation is governed by the interplay between endogenous signaling pathways and exogenous pro-proliferative cues ^[6]^. Beyond serving as energy sources and biosynthetic precursors, it remains unresolved whether metabolic intermediates circulate to mediate inter-organ communication and act as exogenous pro-proliferative signals.

Exercise provides robust cardiovascular protection: long-term aerobic training induces physiological hypertrophy, improves reserve, and limits post-injury remodeling ^[7]^. Some studies suggest exercise can increase cardiomyocyte cell-cycle re-entry, but mechanisms are unclear ^[8]^. Given the profound systemic metabolic effects of exercise, it is plausible that metabolic reprogramming contributes to the regulation of cardiomyocyte proliferation. Yet exercise generally enhances fatty acid oxidation^[8]^, whereas elevated fatty acid oxidation is unfavorable for proliferation ^[4]^—a paradox suggesting that the pro-proliferative effect may not be solely driven by a cardiomyocyte-intrinsic metabolic switch. Exercise-induced inter-organ signals (termed exerkines)—including hormones, metabolites, proteins, and nucleic acids—have broad physiological effects ^[9]^. Exercise-driven shifts in fatty acid metabolism could plausibly trigger system-wide remodeling of the lipidome; yet the dynamics and functions of bioactive lipids within this landscape remain poorly defined. Among these, eicosanoids—bioactive lipids derived from polyunsaturated fatty acids via multiple oxygenases—are particularly compelling candidate exerkines. We previously established a systematic framework for eicosanoid profiling and functional study, and identified several eicosanoids that exert potent effects on cardiomyocyte fate ^[10–12]^, motivating us to ask whether eicosanoids mediate exercise-induced proliferation.

Here, we show that preventive aerobic training reduces infarct-associated injury and accelerates functional recovery. Targeted lipidomics revealed exercise-elevated epoxy eicosanoids, especially epoxyeicosatetraenoic acids (EEQs). Transcriptomics showed enrichment of cell proliferation and cell-cycle programs, with Nuclear Receptor Subfamily 4 Group A Member 1 (NR4A1; TR3/NGFIB/Nur77) as the most prominently upregulated gene ^[13]^. We identify epoxy eicosanoids as exercise-derived mediators that promote cardiomyocyte proliferation and post-infarction repair via an NR4A1-dependent mechanism. Moreover, supplementing eicosapentaenoic acid (EPA), the metabolic precursor of EEQs, boosts EEQ biosynthesis and thereby augments proliferation and cardioprotection, amplifying the benefits of exercise.

## Methods

The authors declare that all supporting data are available within the article and its online supplementary files. The data, all protocols, and study materials are available to other researchers for purposes of reproducing the results. The detailed methods used in this study are provided in the Supplemental Material.

### Statistical Analysis

Data are expressed as mean ± SEM. All statistical analyses were performed using GraphPad Prism 9 software. Differences between the two groups were evaluated using an unpaired, two-tail Studxent’s t-test. For comparison of multiple groups, one-way ANOVA (for one independent variable) followed by Tukey’s post hoc comparison or two-way ANOVA (for two independent variables) followed by Tukey’s post hoc comparison was used. Differences were considered statistically significant if P-value < 0.05.

## RESULTS

### Preventive aerobic training limits infarct injury and accelerates recovery

To test whether prior exercise mitigates post-infarction functional decline, mice underwent 4 weeks of treadmill training or remained sedentary before LAD ligation; cardiac structure and function were assessed thereafter (Figure 1A). Preconditioning with exercise attenuated pathological hypertrophy (Figure 1B) and markedly rescued the infarction-induced reductions in ejection fraction (EF) and fractional shortening (FS), as well as the increases in left ventricular end-systolic and end-diastolic volumes (LVESV, LVEDV) (Figure 1C–1G). Masson’s trichrome staining further showed that prior exercise significantly reduced infarct burden compared with sedentary controls (Figure 1H–1I). To rule out the possibility that improved baseline metabolism accounted for the benefit, we assessed general metabolic parameters. We found that body composition (fat mass and body fat percentage) and glucose homeostasis (determined by glucose tolerance testing) were comparable between exercised and sedentary mice. (Figure S2). Because exercise can remodel lipid metabolism and impact circulating lipids^[14]^, we quantified plasma free fatty acids (FFAs) using LC–MS/MS. Mice completed 4 weeks of aerobic training, and plasma was collected 6 hours after the final session. Although total FFA levels were unaltered, the ratio of polyunsaturated to saturated fatty acids (PUFA/SFA) increased significantly in the exercised group (Figure 1J–1K). Together, these data show that prior aerobic training improves post-infarction structure and function and shifts the composition of circulating FFAs toward a higher PUFA/SFA ratio.

**Figure 1.**
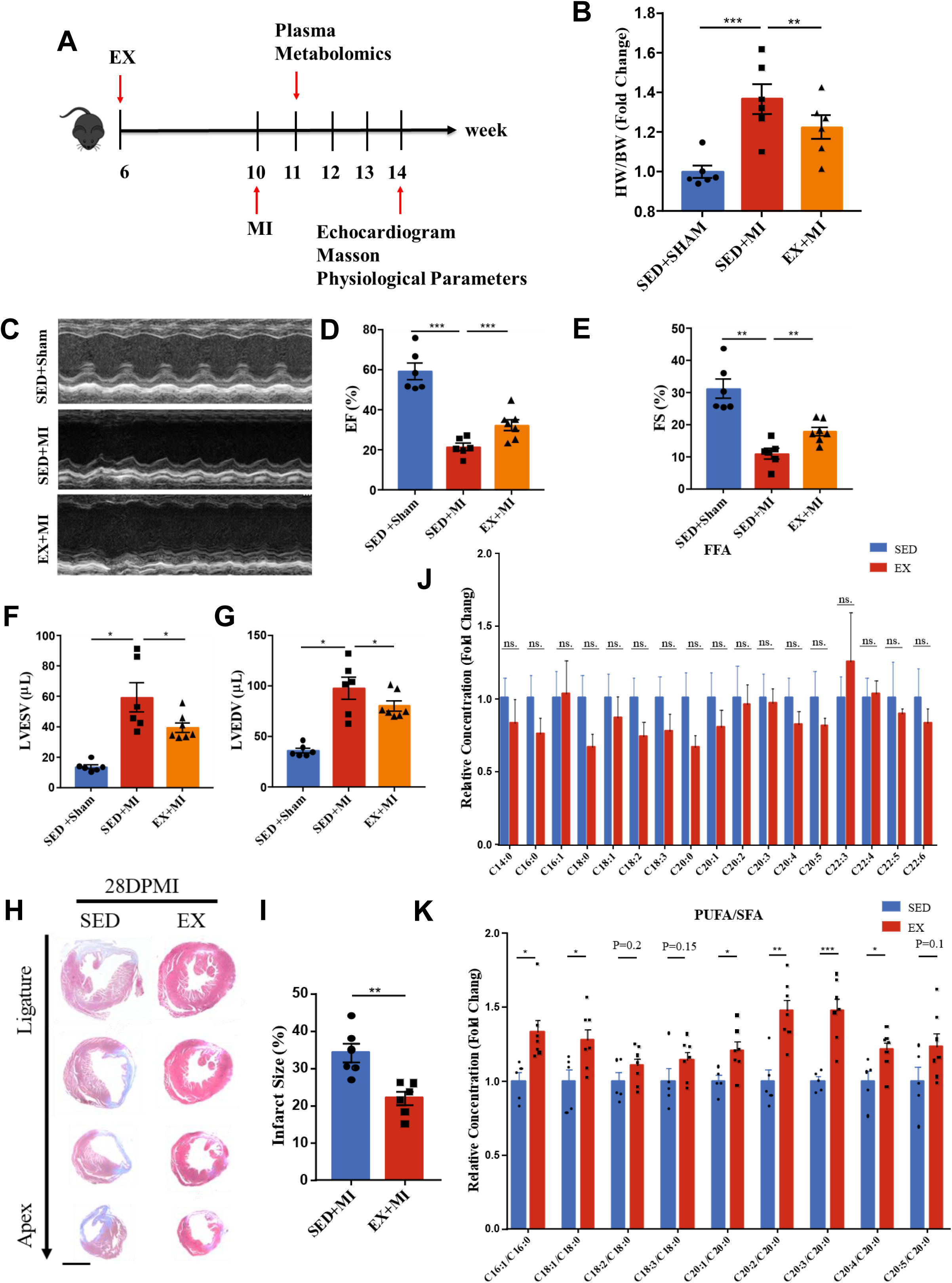
**Pre-exercise confers protective effects against myocardial infarction and increases the PUFA/SFA ratio.** A. Schematic illustration of the experimental timeline for pre-exercise training and subsequent MI induction. B. Heart weight/body weight ratio in sedentary versus pre-exercised mice post-myocardial infarction (n=6). C. Echocardiographic images of sedentary versus pre-exercised mice. D–G. Left ventricular function parameters in sedentary versus pre-exercising mice, including ejection fraction (EF), fractional shortening (FS), left ventricular end-systolic volume (LVESV), and left ventricular end-diastolic volume (LVEDV) (Sham, n=6; SED+MI, n=6; EX+MI, n=7). H. Representative Masson’s trichrome staining of heart sections at 28 days post-MI (scale bar = 1 mm) (n=6). I. Quantification of infarct size based on Masson’s trichrome staining. J. Plasma levels free fatty acid (FFA) levels in sedentary and exercising mice (SED, n=5; EX, n=7). K. Ratio of plasma polyunsaturated fatty acids (PUFA) to saturated fatty acids (SFA) in sedentary and exercising mice (SED, n=5; EX, n=7). Data are presented as mean ± SEM. *P<0.05; **P<0.01; ***P<0.001.

### Exercise remodels the plasma PUFA metabolite profile and increases CYP450-derived epoxy-eicosanoids

In our model, exercise did not significantly lower total FFAs despite a downward trend; however, because SFAs undergo β-oxidation more efficiently than PUFAs, the PUFA/SFA ratio increased after training. Such shifts in plasma fatty acid composition may propagate to tissue lipid distributions, consistent with observational studies reporting long-term exercise–induced increases in membrane PUFA content in skeletal muscle^[15, 16]^. Elevated membrane PUFA content can further reshape eicosanoid metabolism, a diverse class of bioactive lipids that often act as receptor ligands and signal transducers ^[17]^. We therefore hypothesized that specific eicosanoids could serve as “exerkine” mediating post-MI repair. Mice completed 4 weeks of aerobic training, and eicosanoid profiling was performed on plasma collected 7 days after MI. A heatmap revealed global remodeling of the eicosanoid metabolism (Figure 2A). Principal component analysis (PCA) showed that MI induced a shift in eicosanoid profiles, whereas prior exercise redirected the profile along an orthogonal trajectory (Figure 2B). Variable-importance-in-projection (VIP) analysis identified the top 15 contributors to the exercise-associated shift; notably, the top four species—19,20-EDP, 5,6-EET, 16,17-EDP, and 17,18-EEQ—are CYP450-derived epoxy-eicosanoids (Figure 2C). Bar plots of all detected epoxy-eicosanoids demonstrated a coherent increase with exercise preconditioning (Figure 2D–2I). To facilitate understanding of the sources and metabolic fates of epoxy-eicosanoids, we have illustrated the enzymes responsible for their synthesis and degradation. (Figure S1A). Because exercise primarily remodels metabolism in skeletal muscle and liver, we quantified transcripts of CYP450 enzymes known to generate epoxy-eicosanoids in these tissues. Exercise robustly upregulated Cyp2c and Cyp2j family members in skeletal muscle, whereas liver showed no comparable induction. By contrast, the principal catabolic enzyme soluble epoxide hydrolase 2 (Ephx2) was unchanged in both tissues (Figure S3). These data indicate that aerobic training elevates circulating CYP450-pathway metabolites of PUFAs and that their increase is likely driven by enhanced expression of epoxygenases in skeletal muscle rather than changes in epoxy-eicosanoid degradation.

**Figure 2.**
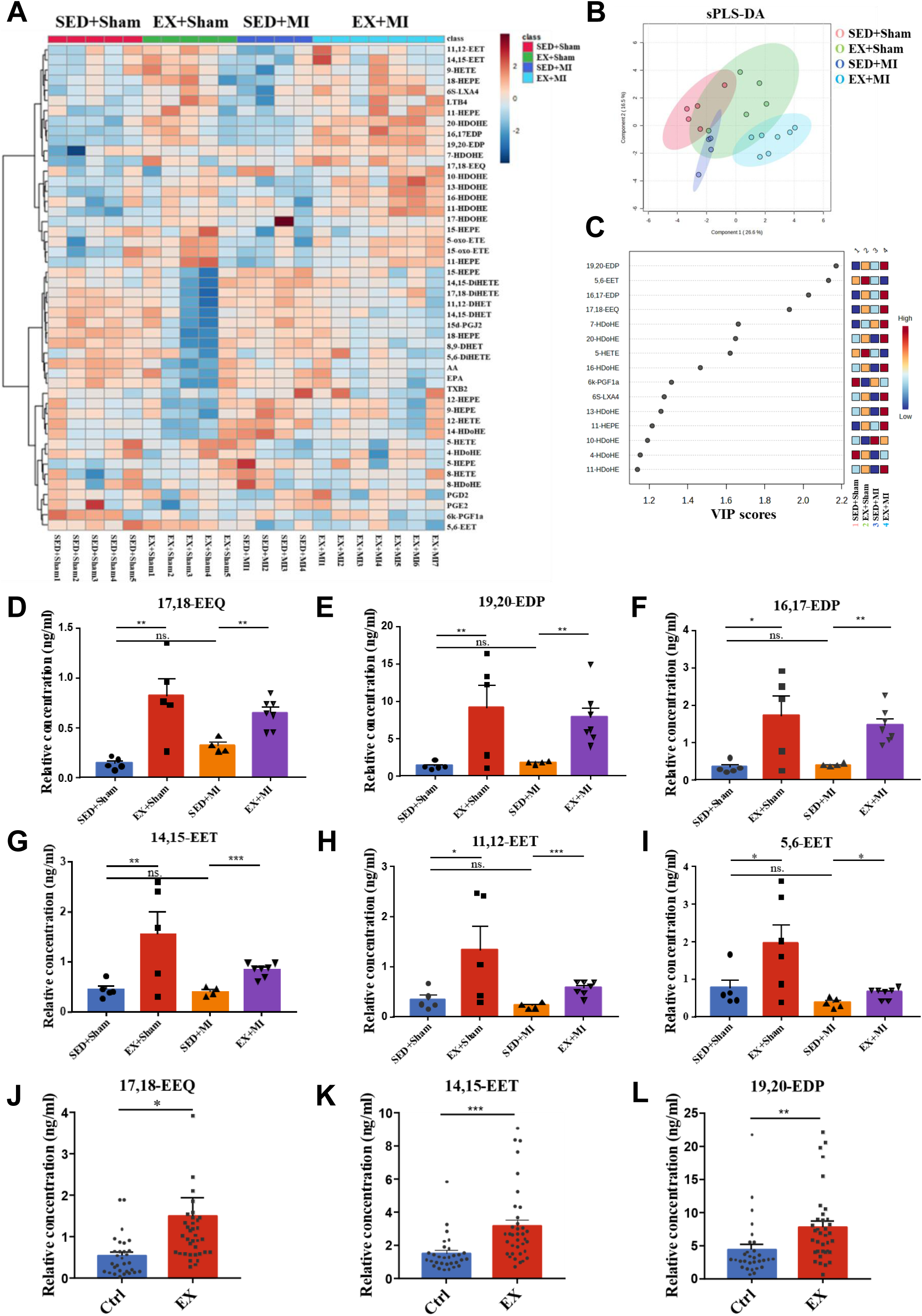
**Eicosanoid profile in sedentary and exercised mice.** A. Heatmap visualization of plasma eicosanoid profiles in sedentary and exercised mice under sham and post-myocardial infarction (MI) conditions (SED, n=5; EX, n=5; SED+MI, n=4; EX+MI, n=7). B. Sparse partial least squares discriminant analysis (sPLS-DA) score plot showing distinct clustering of metabolite profiles between groups. C. Variable Importance in Projection (VIP) scores identifying the top 15 metabolites contributing to group separation. D–I. Plasma concentrations of CYP pathway metabolites 17,18-EEQ, 19,20-EDP, 16,17-EDP, 5,6-EET, 11,12-EET, and 14,15-EET after 28 days of aerobic exercise or sedentary control. J-K. Plasma concentrations of CYP pathway metabolites 17,18-EEQ, 14,15-EET ,19,20-EDP in elite athletes and sedentary controls (age-matched) (Athletes, n=36; Sedentary, n=30). Data are presented as mean ± SEM. *P<0.05; **P<0.01; ***P<0.001.

### Clinical analysis reveals a distinctive plasma epoxy-eicosanoid signature in athletes

To translate our preclinical finding that epoxy-eicosanoids are upregulated by exercise preconditioning, we conducted a clinical study comparing the plasma eicosanoid profiles of elite endurance athletes versus age-matched sedentary controls (Figure S1B). Strikingly, the highest-ranking metabolites were predominantly CYP450-derived epoxy-eicosanoids. Subsequent quantitative analysis focused on these top epoxy-eicosanoid species. Bar plots revealed that the plasma concentrations of six key epoxy-eicosanoids—17,18-EEQ, 14,15-EET, 19,20-EDP, 5,6-EET, 8,9-EET, and 11,12-EET—were all significantly elevated in the Athletes group compared to the Sedentary controls (Figure 1J–1L; Figure S1F–1H). Eicosanoid profiling was performed using LC-MS, and the data were subjected to multivariate analysis. Unsupervised hierarchical clustering, presented as a heatmap, demonstrated a global remodeling of the eicosanoid metabolome, distinguishing the Athletes group from the Sedentary control group (Figure S1C). PCA confirmed a clear separation between the two groups along the principal components, indicating distinct eicosanoid signatures associated with chronic high-level exercise (Figure S1D). To identify the metabolites contributing most to this separation, VIP analysis was performed (Figure S1E). This coherent upregulation of a specific set of epoxy-eicosanoids in highly trained athletes mirrors our prior observation in exercise-preconditioned mice and further supports the association between sustained aerobic training and a marked shift in the plasma epoxy-eicosanoid profile.

### Exercise-associated lipids promote cardiac proliferative programs

To investigate the molecular basis of exercise-enhanced post-MI repair, we performed RNA-seq using tissue collected from the infarct border zone. GO and KEGG enrichment analyses revealed that exercise-regulated genes were prominently enriched in the cell cycle pathway (Figure 3A–3B). GSEA and heatmaps further indicated a positive regulation of cell cycle signatures by aerobic training (Figure 3C–3D). In addition, annotation network illustrated relationships among GO annotations and their enrichment levels; cell cycle–related GO annotations were interrelated and highly enriched (p<10^-10) (Figure S4A–4B). A gene interaction network based on protein–protein interactions (PPI) further demonstrated the correlation structure among differentially expressed genes. The Molecular Complex Detection (MCODE) network analysis component identified topological regions (subnetworks), and genes within the core subnetworks were enriched in GO annotations related to the cell cycle and mitosis (p<10^^-8.6^), further suggesting that exercise preconditioning affects cell cycle/mitosis-related gene networks in the context of MI (Figure S4C). These data suggest that exercise confers cardioprotection, at least in part, by promoting entry into regenerative and proliferative states.

**Figure 3.**
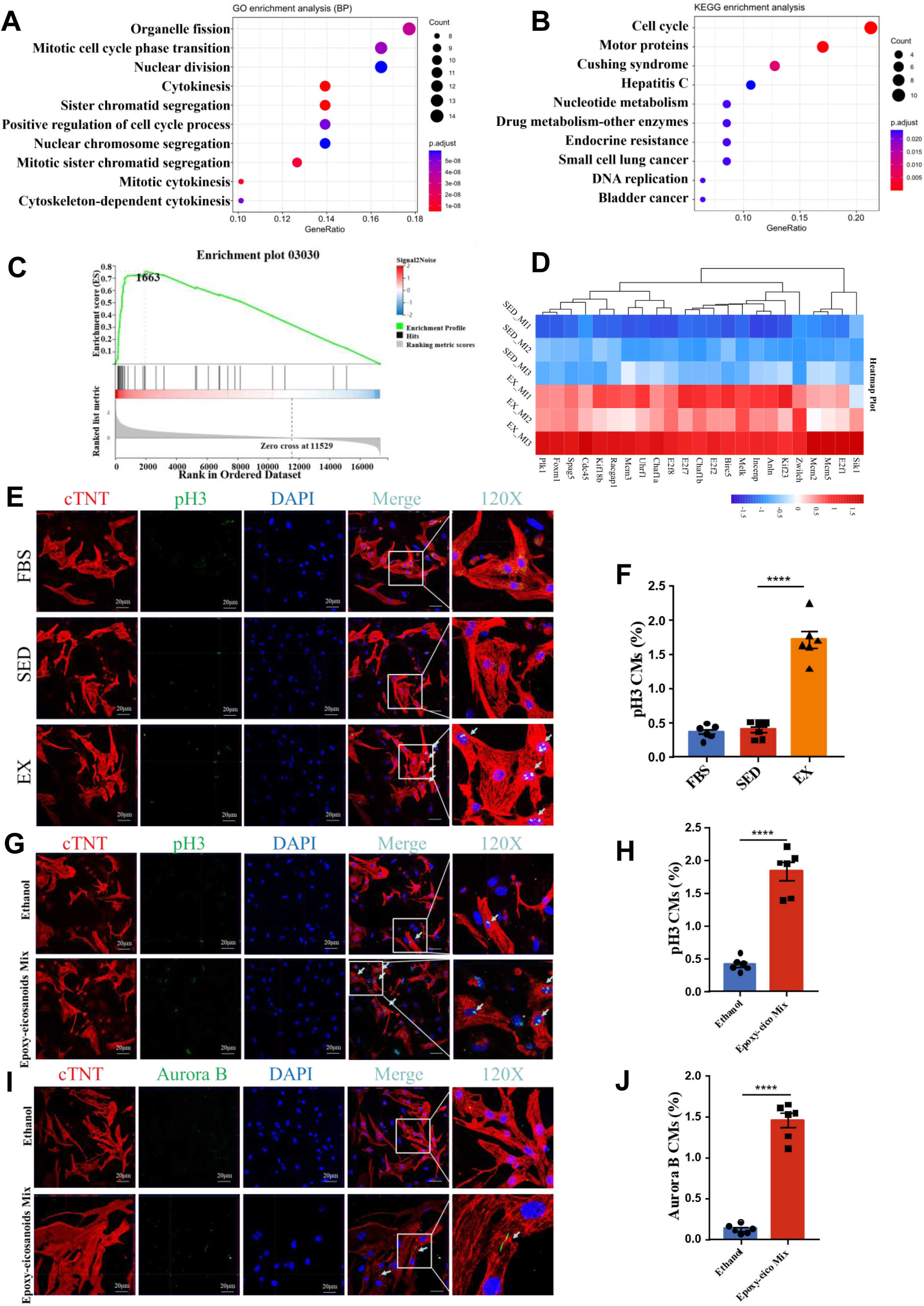
**Exercise-derived bioactive lipid mediators promote cardiomyocyte proliferation post-MI** A–B. Transcriptomic profiling of cardiac tissue from sedentary and exercised mice at 7 days post-MI (n=3 per group). (A) Gene Ontology (GO) and (B) Kyoto Encyclopedia of Genes and Genomes (KEGG) enrichment analyses. Dot color indicates statistical significance (P-value), and dot size represents the number of enriched genes. C. GSEA plot generated from transcriptome sequencing data. Default Signal2Noise algorithm employed. D. Heatmap visualization of differentially expressed genes, with red and blue indicating high and low relative expression levels, respectively. E–F. Proliferative response of neonatal rat cardiomyocytes (NRCMs) to plasma lipid extracts. NRCMs were cultured under hypoxic, serum-free conditions for 24 hours in the presence of lipid mediators extracted from the plasma of sedentary or exercised mice. (E) Representative immunofluorescence images and (F) quantification of phospho-histone H3 (pH3)-positive cells assessed by High-Content Screening (n=6). G–J. Proliferative response of NRCMs to candidate metabolites. NRCMs were treated with a mixture of 17,18-EEQ, 19,20-EDP, and 14,15-EET (or solvent control) for 24 hours under hypoxic, serum-free conditions. (G) Representative images and (H) quantification of pH3 staining. (I) Representative images and (J) quantification of Aurora B kinase staining. Analysis was performed using High-Content Screening (n=6). Data are presented as mean ± SEM. *P < 0.05, **P < 0.01, ***P < 0.001.

We next tested whether plasma fractions (containing exerkines) could influence the cardiomyocyte cell cycle. Primary neonatal rat cardiomyocytes (NRCMs) were used to simulate MI in vitro under hypoxia. Lipid fractions were extracted from plasma of exercised or sedentary mice and applied to NRCMs for 24 h, followed by serum-free hypoxia (1% O2, 5% CO2, 94% N2) for 24 h. Immunofluorescence and immunoblotting showed that lipid extracts from exercised mice increased cardiomyocyte proliferation markers (PH3 and PCNA) relative to sedentary controls (Figure 3E–3F; S5A-5B). Furthermore, we applied a mixture of epoxy-eicosanoid metabolites upregulated by aerobic exercise—17,18-EEQ, 19,20-EDP, and 14,15-EET—to NRCMs, and this mixture significantly increased the expression of Ki67, PH3, and Aurora B after hypoxia (Figure 3G–3J; S5C-5D). Together, these findings identify exercise-induced epoxy-eicosanoids as candidate circulating “exerkines” that promote cardiomyocyte proliferation.

### Exercise-induced epoxy-eicosanoids upregulate NR4A1 to drive cardiomyocyte proliferation

To identify mediators of the pro-proliferative effect, we screened exercise-regulated genes in post-MI hearts and found NR4A1 as the most prominently upregulated candidate under stringent criteria (Q<0.1, log2FC>2; Figure 4A). We validated increased NR4A1 expression in exercised post-MI hearts at both the protein level by immunoblotting (Figure 4B-4C) and the transcript level by qPCR (Figure 4D). To test whether epoxy-eicosanoids act via NR4A1, primary NRCMs were exposed to epoxy-eicosanoids under hypoxia (1 μM, 24 h). Each epoxy-eicosanoid elevated NR4A1 mRNA and protein level, with 17,18-EEQ producing the strongest induction (Figure 4E–4F; Figure S6A). Consistently, 17,18-EEQ most robustly increased cardiomyocyte proliferation markers (Ki67, PH3, and Aurora B) under hypoxia compared with vehicle (Figure 4G–4J; Figure S6B-6C). Together, these results identify NR4A1 as an exercise-responsive cardiac factor and suggest that 17,18-EEQ promotes the expression of NR4A1.

**Figure 4.**
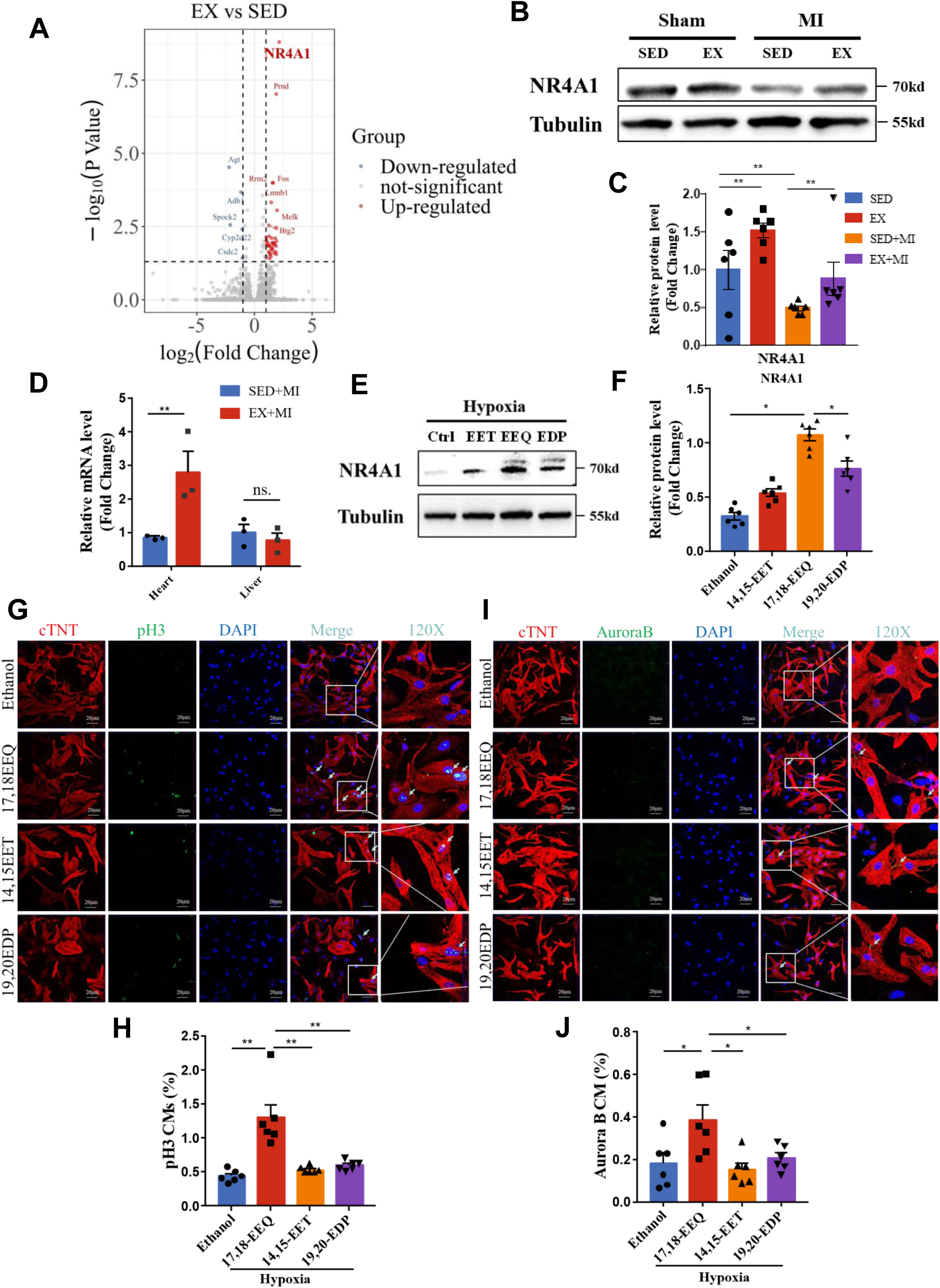
**17,18-EEQ upregulates exercise -responsive protein NR4A1 and promotes cardiomyocyte proliferation.** A. Volcano plot of differentially expressed genes identified by RNA-seq analysis of cardiac tissue from sedentary and exercised mice at 7 days post-MI (n=3). Red and blue indicate high and low relative expression levels, respectively. B-C. Protein expression and statistical analysis of NR4A1 in the myocardial infarct region and liver tissue of mice undergoing 28 days of aerobic exercise (n=6). D. mRNA expression levels of Nr4a1 in the myocardial infarct region and liver tissue of sedentary versus exercised mice (n=3). E-F. Induction of NR4A1 protein by specific metabolites. NRCMs were treated with 14,15-EET, 17,18-EEQ, or 19,20-EDP for 24 hours under hypoxic, serum-free conditions. (E) Representative Western blots and (F) quantification of NR4A1 expression (n=6). G-J. Effect of specific metabolites on cardiomyocyte proliferation. NRCMs were treated with 14,15-EET, 17,18-EEQ, or 19,20-EDP for 24 hours under hypoxic, serum-free conditions. (G) Representative images and (H) quantification of pH3 staining. (I) Representative images and (J) quantification of Aurora B staining. Analysis was performed using High-Content Screening (n=6). Data are presented as mean ± SEM. *P < 0.05, **P < 0.01.

### 17,18-EEQ activates the cAMP–PKA–CREB–NR4A1 axis to promote cardiac proliferation and repair

Given that NR4A1 is sensitive to upstream phosphorylation signals^[18]^, we performed phosphoproteomic profiling of hypoxic NRCMs pretreated with 17,18-EEQ. Gene Ontology (GO) enrichment analysis of the differentially phosphorylated proteins highlighted significant associations with PKA regulatory subunit binding (Molecular Function, MF) and cell-cycle pathways (Biological Process, BP) (Figure 5A–5B). Specifically, within the phosphoproteomic dataset, we observed that 17,18-EEQ treatment markedly increased the phosphorylation of the canonical PKA substrate, CREB (Figure 5C). To validate this finding, we performed time-course Western blot analyses under hypoxia, which confirmed that 17,18-EEQ elevated p-CREB levels as early as 1 h post-treatment (Figure 5D–5E). This activation was further corroborated by immunofluorescence staining (Figure 5F–5G). Consistent with reports that the cAMP–PKA–CREB axis regulates NR4A1 expression^[19, 20]^, ELISA assays confirmed that 17,18-EEQ increased intracellular cAMP levels (Figure 5J). To determine the functional necessity of this signaling cascade, we pharmacologically inhibited distinct nodes using the adenylyl cyclase inhibitor SQ22536 and the PKA inhibitor H89. Notably, both inhibitors abolished the 17,18-EEQ–induced increases in Ki67 and PH3 (Figure 5K–5N; Figure S7), and H89 significantly attenuated NR4A1 upregulation (Figure 5H–5I). Collectively, these findings demonstrate that 17,18-EEQ drives cardiomyocyte proliferation by activating the cAMP–PKA–CREB axis, which subsequently induces NR4A1 expression.

**Figure 5.**
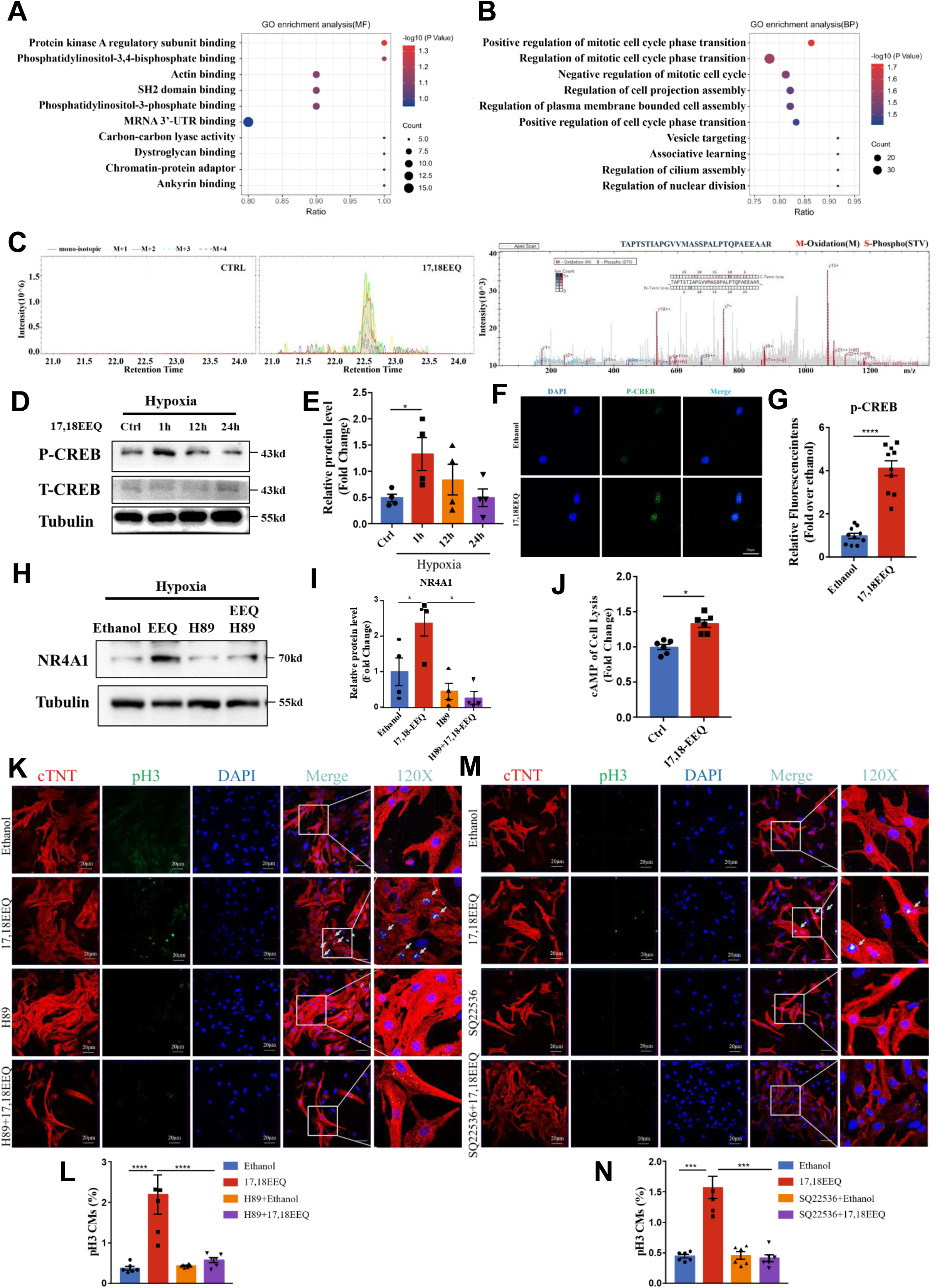
**17,18-EEQ promotes cardiomyocyte proliferation via the cAMP-PKA-CREB pathway in vitro.** A–B. GO analysis of protein phosphorylation profiles following 17,18-EEQ stimulation of NRCMs under hypoxic, serum-free conditions. C. Extracted ion chromatogram (XIC) and b, y ion pair diagram of the CREB phosphorylated peptide sequence following 17,18-EEQ stimulation of NRCMs under hypoxic, serum-free conditions. D-E. Time-course analysis of CREB phosphorylation. NRCMs were treated with 17,18-EEQ for 1, 12, and 24 hours under hypoxic, serum-free conditions. (D) Representative Western blots and (E) quantification of the p-CREB to total CREB (t-CREB) ratio (n=4). F-G. Nuclear translocation of p-CREB. (F) Representative immunofluorescence images and (G) quantification of p-CREB intensity in NRCMs treated with 17,18-EEQ (n=10). H-I. PKA-dependent regulation of NR4A1. NRCMs were treated with 17,18-EEQ in the presence or absence of the PKA inhibitor H89. (H) Representative Western blots and (I) quantification of NR4A1 protein levels (n=4). J. cAMP levels in NRCMs stimulated with 14,15-EET and 17,18-EEQ under hypoxic, serum-free conditions (n=6). K-N. Requirement of cAMP/PKA signaling for proliferation. NRCMs were treated with 17,18-EEQ in the presence of the PKA inhibitor H89 (K–L) or the adenylyl cyclase inhibitor SQ22536 (M–N). (K, M) Representative immunofluorescence images and (L, N) quantification of pH3-positive cells assessed by High-Content Screening (n=6). Data are presented as mean ± SEM. *P < 0.05, ***P < 0.001.

### In Vivo Validation of NR4A1 as a Mediator of Exercise-Induced Cardiomyocyte Proliferation and Post-MI Repair

To test whether exercise confers cardioprotection via NR4A1, we systemically delivered AAV9-shNR4A1 (control: AAV9-empty) to knock down NR4A1 in mice (Figure 6A). AAV9-shNR4A1 achieved robust cardiac knockdown of NR4A1 at both mRNA and protein levels (Figure 6B–6C). NR4A1 depletion abrogated the exercise preconditioning benefit on cardiac function after MI, reversing improvements observed in echocardiographic indices (Figure 6D–6F). Histopathology further showed that NR4A1 knockdown abolished the exercise-induced reduction in fibrosis area (Figure 6G–6H). Concordantly, the exercise-driven increases in the cardiomyocyte proliferation markers Ki67 and pH3 were also reversed when NR4A1 was depleted (Figure 6I–6J; Figure S8). These data demonstrate that NR4A1 is required for the pro-proliferative and cardioprotective effects of aerobic exercise after MI.

**Figure 6.**
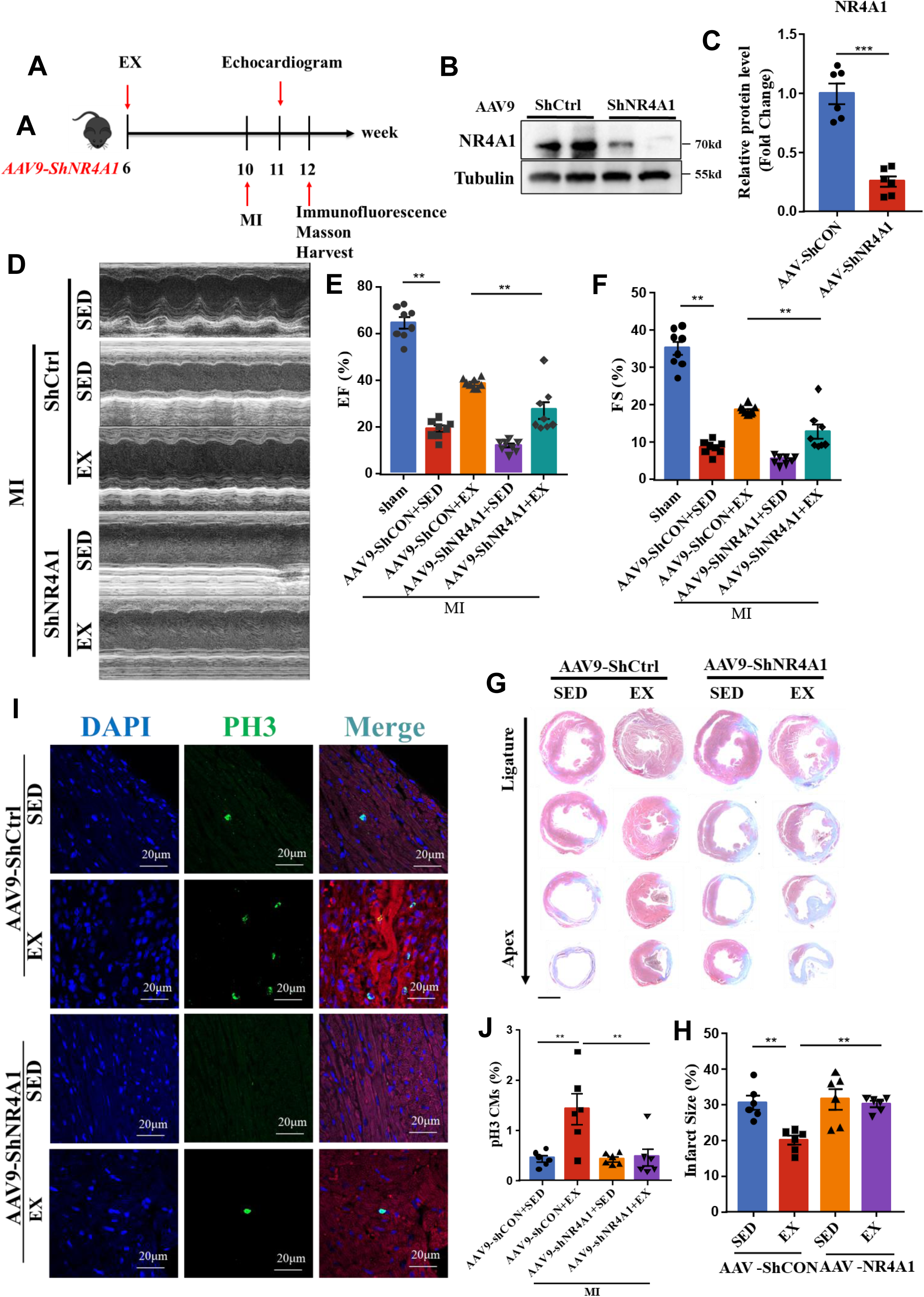
**Cardiac-specific knockdown of NR4A1 abolishes exercise-induced cardioprotection post-MI.** A. Schematic illustration of the experimental protocol involving AAV9-mediated knockdown, exercise training, and MI induction. B-C. Verification of knockdown efficiency. (B) Representative Western blots and (C) quantification of NR4A1 protein levels in heart tissue following AAV9-shNR4A1 (or control AAV9) injection (n=6). D. Representative echocardiograms of exercise and sedentary mice after myocardial infarction following AAV9-NR4A1 knockdown. E–F. Echocardiographic assessment of cardiac function. Left ventricular ejection fraction (LVEF) (E) and fractional shortening (LVFS) (F) in the indicated groups (n=8). G–H. Assessment of infarct size. (G) Representative Masson’s trichrome staining of heart sections and (H) quantification of the fibrotic area (n=6). I–J. Evaluation of cardiomyocyte proliferation in vivo. (I) Representative immunofluorescence images of Ki67-positive cells in the border zone of the infarct and (J) quantification of Ki67-positive cardiomyocytes (n=6). Data are presented as mean ± SEM. *P < 0.05, **P < 0.01.

### In vivo validation that the 17,18-EEQ–PKA–CREB–NR4A1 axis promotes cardiomyocyte proliferation and post-MI repair

To assess the cardioprotective effect of the exercise-derived lipid 17,18-EEQ in vivo, we implanted subcutaneous ALZET osmotic pumps to deliver 17,18-EEQ for four weeks and performed MI in week 3 (Figure 7A). Pump delivery increased circulate 17,18-EEQ levels (Figure 7B), reduced fibrosis area (Figure 7C–7D), and improved cardiac function after acute MI (Figure 7E–7F). Consistently, 17,18-EEQ elevated cardiomyocyte proliferation markers Ki67, pH3, and Aurora B in the heart (Figure 7G–7L). We next validated the mechanism in vivo. Mice receiving 17,18-EEQ pumps were co-treated with inhibitors targeting distinct nodes of the pathway—SQ22536 (adenylyl cyclase/cAMP), H89 (PKA), and the CREB phosphorylation inhibitor 666-15—via intraperitoneal injection (Figure 8A–8B). Compared with 17,18-EEQ alone, each inhibitor increased fibrosis area (Figure 8C–8D), abolished the post-MI functional benefits (Figure 8E–8F), and reversed the increases in Ki67, pH3, and Aurora B (Figure 8G–8L). In summary, these data demonstrate that 17,18-EEQ acts through the cAMP–PKA–CREB pathway to regulate downstream NR4A1 expression and cardiomyocyte proliferation.

**Figure 7.**
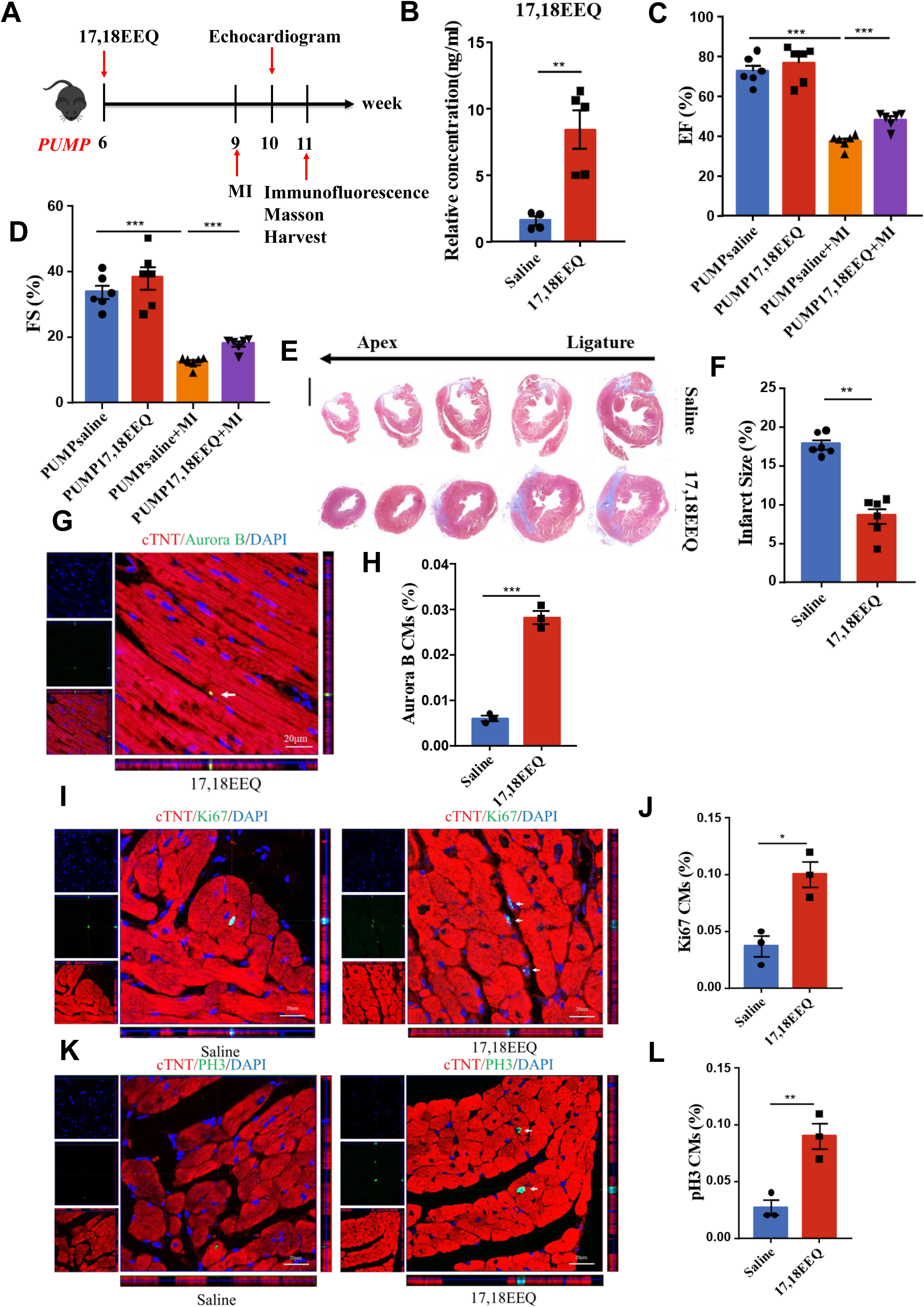
**Administration of 17,18-EEQ promotes cardiomyocyte proliferation and improves cardiac repair post-MI.** A. Schematic illustration of the experimental protocol: subcutaneous osmotic pump implantation for 17,18-EEQ (or saline) delivery followed by MI induction. B. Plasma 17,18-EEQ concentrations after subcutaneous pump delivery of saline versus 17,18-EEQ (n=3). C–D. Assessment of infarct size. (C) Representative Masson’s trichrome staining of heart sections and (D) quantification of the fibrotic area (n=6). E–F. Echocardiographic assessment of cardiac function. Left ventricular ejection fraction (LVEF) (E) and fractional shortening (LVFS) (F) in mice treated with saline or 17,18-EEQ (n=6). G-L. Evaluation of cardiomyocyte proliferation in vivo. Representative immunofluorescence images and quantification of (G–H) Aurora B, (I–J) Ki67, and (K–L) pH3 positive cells in the border zone of the infarct. Analysis was performed using High-Content Screening (n=3). Data are presented as mean ± SEM. *P < 0.05, **P < 0.01; **P < 0.001.

**Figure 8.**
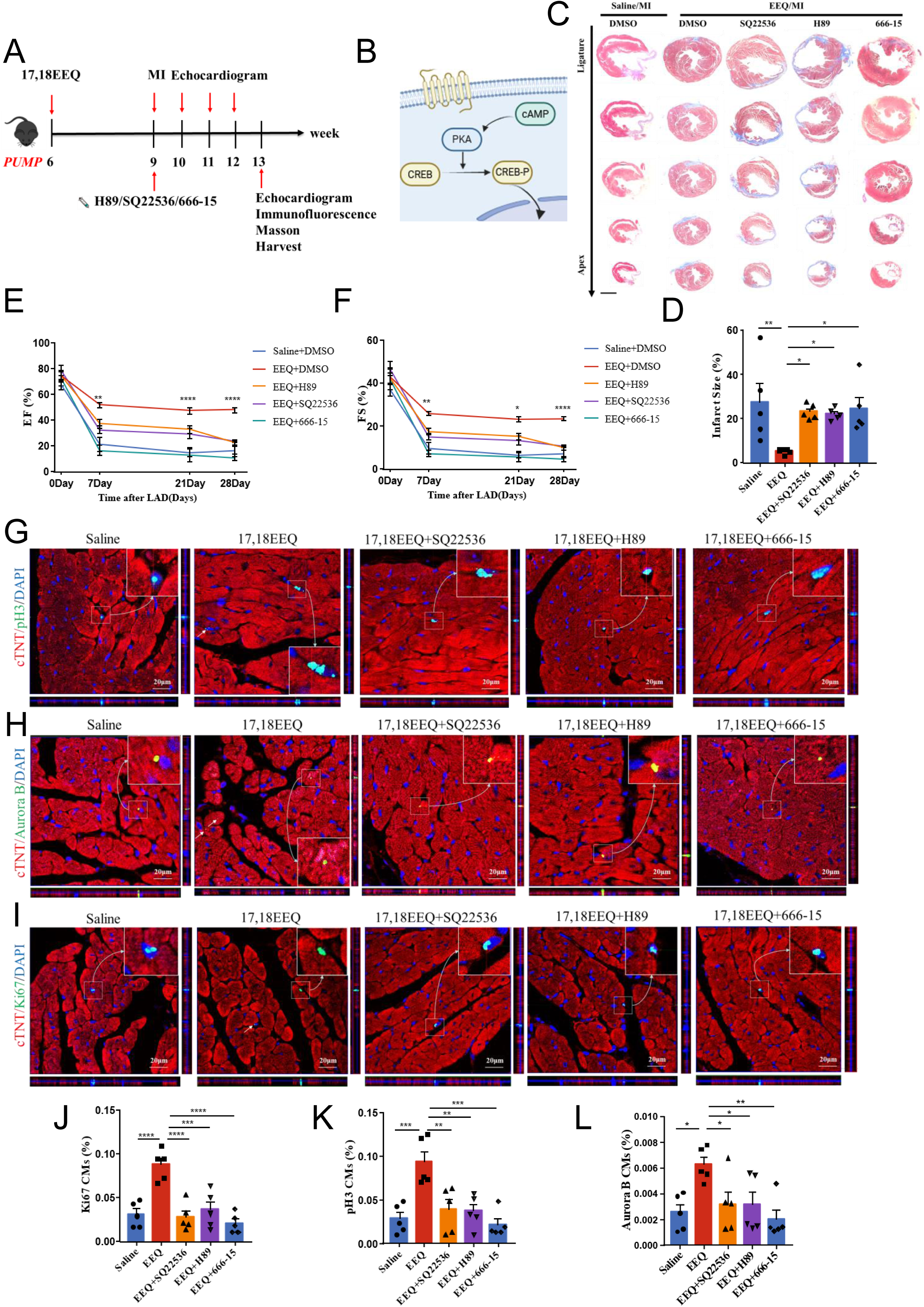
**17,18-EEQ promotes cardiomyocyte proliferation and cardiac repair in vivo via the cAMP-PKA-CREB pathway.** A. Schematic illustration of the experimental design involving continuous subcutaneous delivery of 17,18-EEQ alone or in combination with specific inhibitors (H89, SQ22536, or 666-15) post-MI. B. Schematic of the cAMP-PKA-CREB signalling pathway. C-D. Assessment of infarct size. (C) Representative Masson’s trichrome staining and (D) quantification of the fibrotic area in hearts treated with 17,18-EEQ in the presence or absence of the indicated inhibitors (n=5). E-F, Echocardiographic assessment of cardiac function. Left ventricular ejection fraction (LVEF) (E) and fractional shortening (LVFS) (F) in the indicated treatment groups (n=5). G-L. Evaluation of cardiomyocyte proliferation in vivo. Representative immunofluorescence images and quantification of (G–H) Ki67, (I–J) pH3, and (K–L) Aurora B positive cells in the border zone. Mice were treated with 17,18-EEQ with or without the indicated inhibitors. Analysis was performed using High-Content Screening (n=5). Data are presented as mean ± SEM. *P < 0.05, **P < 0.01, ***P < 0.001, ****P < 0.0001.

### Omega-3 supplementation amplifies the cardioprotective benefit of aerobic exercise after MI

Finally, we investigated whether other interventions that increase EEQs could augment the cardioprotective effect of exercise. The metabolic precursor of 17,18-EEQ is EPA, an ω-3 PUFA. Because mammals lack ω-desaturase, they cannot de novo synthesize ω-3 PUFAs, and thus ω-3 PUFAs are mainly obtained from diet; ω-3 PUFAs are recommended by the FDA as cardioprotective dietary components^[21]^. We provided mice with omega-3-enriched chow during aerobic exercise training (Figure S9A). Compared with either exercise or omega-3 alone, the combination further improved post-MI cardiac function and reduced fibrosis area (Figure S9B–9E). Metabolomics revealed distinct metabolic profiles in exercised and omega-3-fed mice versus sedentary controls (Figure S10A–10C), and their combination further increased in vivo 17,18-EEQ levels (Figure S10D), with 17,18-EEQ ranking as the top contributor to the metabolic signature (Figure S10E).

## DISCUSSION

In this study, we demonstrated the robust cardioprotective effects of exercise preconditioning. By focusing on exercise-driven metabolic modulation, we utilized metabolomics to identify the bioactive lipid 17,18-EEQ as a critical exercise-derived mediator that directly confers protection against myocardial infarction injury by promoting cardiomyocyte proliferation. Complementary transcriptomic analysis pinpointed NR4A1 as the central effector mediating the benefits of both exercise and 17,18-EEQ. Furthermore, phosphoproteomic profiling elucidated a cAMP/PKA/CREB signaling cascade connecting 17,18-EEQ to NR4A1, thereby delineating a coherent mechanistic axis. These findings not only expand our understanding of exercise-induced cardiovascular protection but also unveil novel therapeutic targets and actionable strategies for MI prevention and repair. Additionally, this study offers a viable approach to stimulate cardiomyocyte proliferation in the adult heart.

### Prevention of myocardial infarction injury by exercise and the discovery of a novel exerkine

Ischemic heart disease remains the leading cause of death worldwide, underscoring the need for safe and effective strategies for its prevention and treatment^[22, 23]^. Clinical studies indicate that exercise confers cardiovascular benefits^[24]^. Notably, individuals who engage in long-term exercise exhibit better outcomes even when MI occurs, compared with sedentary counterparts ^[25–27]^. In our study, animal experiments demonstrated that pre-exercise facilitates post-MI repair, indicating that exercise is indeed an effective preventive strategy against ischemic heart disease. Several studies have also shown that early post-MI exercise can improve ventricular remodeling, exercise tolerance, and autonomic balance, thereby supporting exercise as a rehabilitation strategy for MI ^[28]^. However, post-MI exercise may increase cardiac load and the risk of complications in some patients ^[29]^. Therefore, compared with post-MI rehabilitation, we advocate moderate-intensity preventive exercise for individuals at risk of ACS, consistent with the AHA’s perspective^[24]^.

Regular, long-term preventive exercise reduces the incidence and mortality of ischemic heart disease through multiple mechanisms, including improving blood pressure, lipids, and insulin sensitivity; reducing inflammation; enhancing endothelial function; and promoting coronary microcirculation and collateral formation ^[30]^. In our animal models, we found that the transcriptomic impact of exercise was characterized by the promotion of cell-cycle entry into mitosis, which complements the existing mechanisms of exercise-induced cardioprotection. Furthermore, the field increasingly recognizes that exercise exerts systemic effects via the release of ’exerkines’—signaling molecules that mediate organ-to-organ communication through the circulation. We identified 17,18-EEQ as a novel exercise-derived bioactive lipid that fulfills the criteria of an exerkine: it is synthesized in peripheral tissues and acts distally on the heart. Given that skeletal muscle is now widely appreciated as an endocrine organ, we investigated its potential role as a source of 17,18-EEQ. We observed a marked upregulation of transcripts encoding 17,18-EEQ biosynthetic enzymes in skeletal muscle following exercise. This aligns with prior findings that exercise-induced lipid remodeling increases the proportion of PUFAs in muscle membranes, thereby providing an abundant substrate pool for 17,18-EEQ production. Collectively, these lines of evidence suggest that post-exercise skeletal muscle acquires the necessary metabolic machinery and substrates to generate 17,18-EEQ, likely serving as a primary source of its systemic release. The cardioprotective action of 17,18-EEQ thus underscores the critical importance of skeletal muscle–myocardium crosstalk in exercise physiology.

### Significance of cardiomyocyte proliferation

Ischemic heart disease is fundamentally driven by progressive cardiomyocyte loss that overwhelms the limited regenerative capacity of the adult heart. While current assistive therapies mitigate secondary damage, they fail to replenish lost cardiomyocytes, inevitably fueling adverse remodeling and heart failure with reduced ejection fraction (HFrEF). In contrast, additive strategies that activate endogenous cardiomyocyte proliferation can directly restore contractile mass and enhance MI repair outcomes beyond current therapeutic limits. Through multi-omics analyses, we revealed that the cardioprotective efficacy of exercise is centrally linked to heightened cardiac cell-cycle activity. Specifically, we delineate a 17,18-EEQ–NR4A1 signaling axis that promotes cardiomyocyte proliferation in the context of MI. Activation of this axis significantly increased the number of Ki67+ and pH3+ cardiomyocytes in adult hearts, indicating enhanced mitotic entry, and elevated the proportion of Aurora B+ cells, confirming progression into substantive cytokinesis. These cellular events translated into tissue-level benefits, including reduced scar size and restored ejection fraction. Collectively, these data demonstrate that activating this axis confers MI protection by promoting cardiomyocyte proliferation with distinct ’regeneration-like’ features, thereby providing a substantive addition to the theoretical framework of cardiac regeneration.

Current views posit that metabolic reprogramming is among the most promising strategies to enhance cardiomyocyte proliferation and promote cardiac regeneration. Shifting energy production from FAO to glycolysis facilitates cardiomyocyte cell-cycle re-entry by supplying biosynthetic precursors and remodeling the epigenetic landscape of proliferation-related genes. However, a paradox remains: during MI, cardiomyocytes are hypoxic and glycolysis is passively upregulated, yet robust proliferation fails to occur, implying the existence of additional checkpoints or missing stimuli. While long-term exercise generally enhances FAO—a metabolic state not intrinsically favorable to a glycolytic shift—we observed significant exercise-induced lipidome remodeling. PUFAs contribute relatively little to β-oxidation yet engage distinct metabolic routes that generate eicosanoid-like mediators with potent bioactivity. Here, we identified the epoxy-eicosanoid 17,18-EEQ as a metabolic driver of proliferation that operates distinct from, yet complementary to, the classical “glycolytic switch”. Intriguingly, 17,18-EEQ promoted proliferation exclusively under hypoxia, not normoxia. This suggests that the passively enhanced glycolysis inherent to the hypoxic MI microenvironment may provide the necessary metabolic prerequisites for proliferation competence. We therefore posit that the proliferation-boosting effect of exercise-derived 17,18-EEQ is contingent on the hypoxic milieu intrinsic to MI.

### Significance and novelty of the 17,18-EEQ–NR4A1 signaling axis

We identify a previously unrecognized regulatory effect of 17,18-EEQ on NR4A1. Prior work showed that 17,18-EEQ suppresses endothelial inflammation and exerts anti-atherosclerotic activity ^[31]^, suggesting that its anti-inflammatory properties may partly contribute to post-MI repair. Notably, fine-tuned immunomodulation itself can facilitate cardiomyocyte proliferation ^[32, 33]^. Mechanistically, the pathway delineated here is distinct from earlier observations: pharmacological reversal of 17,18-EEQ’s effects indicates that its benefit in MI repair more likely arises from directly promoting cardiomyocyte proliferation. Taken together with our previous findings in atherosclerosis, these data position 17,18-EEQ as a potential ’full-chain’ therapeutic target across the spectrum of atherosclerotic cardiovascular disease (ASCVD).

NR4A1, an orphan nuclear receptor encoded by an immediate early gene, serves as a rapid responder to external stimuli independent of de novo protein synthesis, initiating downstream cascades with profound regulatory impact^[34, 35]^. In our study, exercise upregulated NR4A1 at both transcript and protein levels in the post-MI myocardium, concomitant with elevated cell-cycle markers Ki67 and pH3. Crucially, AAV9-mediated knockdown of NR4A1 abolished the exercise-induced improvements in cardiac function, fibrosis reduction, and cardiomyocyte proliferation, establishing NR4A1 as an indispensable mediator.

Interestingly, NR4A1 is known to regulate proliferation bidirectionally, contingent on cellular context^[36, 37]^. For instance, in liver regeneration, an NR4A1–YAP feedback loop coordinates proliferation and apoptosis by promoting YAP ubiquitination ^[38]^. Given that YAP is a potent driver of myocardial regeneration^[39]^, these parallels suggest that cardiac regeneration may rely on similar coordinated pathway interactions. Whether exercise-induced NR4A1 modulation intersects with YAP signaling in the post-MI heart needs further investigation.

To elucidate the signal transduction linking 17,18-EEQ to NR4A1, our phosphoproteomic analysis implicated the cAMP–PKA–CREB axis. This aligns with prior reports in neurons, where the cAMP–PKA–CREB pathway upregulates NR4A1 to confer neuroprotection ^[40]^. Our findings confirm that this signaling paradigm is conserved in cardiomyocytes, serving as the bridge between 17,18-EEQ stimulation and NR4A1-driven regenerative responses.

### Translational relevance

We identified 17,18-EEQ as a critical exercise-induced mediator that confers robust protection against MI. Given that its metabolic biosynthetic pathway is well-defined, strategies targeting this axis hold significant clinical potential. EPA, the specific substrate for 17,18-EEQ, is an omega-3 PUFA with low endogenous abundance in mammals and must therefore be obtained primarily through diet. In this study, we demonstrated that dietary omega-3 PUFA supplementation synergistically augments the cardioprotective efficacy of exercise post-MI. This finding not only substantiates the pivotal role of 17,18-EEQ but also establishes a feasible, pathway-directed intervention strategy.

Notably, while omega-3 PUFAs are widely regarded as cardioprotective, recent clinical trials have yielded inconsistent results regarding their ability to improve outcomes in MI patients ^[41, 42]^. Our findings suggest that this discrepancy may stem from an insufficient metabolic conversion of precursor omega-3 PUFAs into potent bioactive lipid mediators, such as 17,18-EEQ. We propose that exercise acts as a necessary catalyst to mobilize the enzymatic machinery required for this conversion, thereby unlocking the full therapeutic potential of omega-3 supplementation.

## CONCLUSION

This study identifies a lipid–signaling axis that links exercise to cardiac proliferation. Preventive aerobic training increases CYP450-derived epoxy-eicosanoids, particularly 17,18-EEQ, which activate a cAMP–PKA–CREB pathway to upregulate NR4A1, thereby driving cardiomyocyte proliferation and limiting infarct injury. Genetic and pharmacologic perturbations demonstrate that NR4A1 is required and that the pathway is necessary for both proliferation and functional recovery. In vivo delivery of 17,18-EEQ recapitulates the benefits of exercise, while omega-3 supplementation enhances endogenous EEQ biosynthesis and further augments protection. These findings provide a mechanistic basis for exercise-induced cardioprotection and nominate the 17,18-EEQ–NR4A1 axis as a practical target for MI prevention and therapy.

## ARTICLE INFORMATION

### Disclousures

J.W., X.Z., Y.Z. have filed patent applications related to cardiac repair by 17,18EEQ.

## Supplemental Materials

Extended Methods

Figure S1–S9

Tables S1–S2

## Supplemental Material

### Experimental Model and Subject Details

#### Myocardial Infarction Surgery

All animal experiments were conducted in accordance with relevant ethical guidelines and were approved by the Animal Ethics and Use Committee of Tianjin Medical University (Tianjin, China). Six-week-old male C57BL/6J mice weighing approximately 20 g were used. Mice were fixed in a supine position on a constant temperature plate (37°C), and the precordial area was disinfected with povidone-iodine. Anesthesia was induced and maintained by inhalation of 3% isoflurane (flow rate 1 L/min). Under a surgical lamp, the point of maximal cardiac impulse was identified, and a small incision was made with sterile scissors. A purse-string suture (4-0) was placed around the opening. The precordial muscles were bluntly dissected using curved forceps. The pleura and intercostal muscles were carefully spread at the fourth intercostal space with blunt-tipped curved forceps. The heart was gently extruded from the thoracic cavity by applying light pressure to the chest and sub-diaphragmatic area, avoiding excessive manipulation to prevent severe pneumothorax. The heart was positioned at the incision site. A 6-0 suture was used to ligate the left anterior descending (LAD) coronary artery. The ligation was secured with two surgical knots, with a needle depth just sufficient to be visible to avoid penetrating the cardiac chamber and causing major hemorrhage, and a width of approximately 1 mm to ensure complete vessel occlusion. For the sham group, the suture was passed under the artery but not tied. After ligation, the excess suture was trimmed, the heart was immediately returned to the thoracic cavity, and air was expelled by gently pushing the diaphragm upwards. The purse-string suture was tightened, the skin was sutured, and the mouse was allowed to recover with supplemental oxygen and warmth before being returned to its cage.

#### Immunocytochemistry

Cells were seeded onto glass coverslips placed in 6-well plates. After 24 hours of attachment, cells were treated according to the experimental design. Cells were fixed with 1 mL of 4% paraformaldehyde (PFA) for 15-20 minutes at room temperature. Subsequently, cells were permeabilized with 1 mL of 0.5% Triton X-100 in PBS for 15 minutes. Non-specific binding was blocked with 500 μL of goat serum blocking buffer for 1 hour. Between each step, cells were washed three times with PBS for 5 minutes each. Primary antibodies were applied and incubated overnight at 4°C. After washing, secondary antibodies were applied and incubated for 1 hour at room temperature in the dark. Finally, coverslips were mounted using a mounting medium containing DAPI.

#### Echocardiography

Six-week-old male mice (25 g) were depilated on the anterior chest area. Cardiac function and ventricular structure were assessed using a Vevo 2100 imaging system. Mice were anesthetized with 3% isoflurane (1 L/min) and fixed on a 37°C platform. A 30 MHz ultrasound probe was used to examine the heart. Ultrasound gel was applied to the left chest. Clear long-axis views of the heart were obtained in B-mode. The probe was held steady, switched to M-mode, and the images were acquired. Parameters from four consecutive long-axis cardiac cycles were averaged. The left ventricular trace was used to measure left ventricular internal diameter at end-diastole (LVIDd) and end-systole (LVIDs), left ventricular end-diastolic volume (LVEDV), left ventricular end-systolic volume (LVESV), left ventricular ejection fraction (LVEF), and fractional shortening (FS).

#### Aerobic Treadmill Training Protocol

Mice underwent aerobic exercise for 5 consecutive days, followed by 2 days of rest each week. To accommodate their nocturnal activity patterns and avoid disrupting their circadian rhythm, training sessions were conducted from 21:00 to 22:00. The treadmill parameters were set as follows: speed 15 m/min, duration 60 min, incline 15°, and a mild electric stimulus (3 Hz, 15 shock limit) at the rear of the lane to encourage running. Mice in the exercise group were placed on the moving treadmill. Mice in the sedentary group were placed on the treadmill, but the belt was not activated. All other housing and handling conditions were identical for both groups. Mice that repeatedly touched the electric grid were removed from the session. After 60 minutes, the treadmill was stopped, and mice were returned to their home cages.

#### Western Blot

Add protein samples sequentially into the gel wells according to the experimental design. Connect the Bio-Rad electrophoresis unit to the power supply, maintaining a voltage of 70V in the upper concentrating gel section and 120V in the lower separating gel section. Once the bromophenol blue protein band has migrated to the bottom of the lower separating gel, promptly switch off the power supply. For film transfer, connect the power supply to the Bio-Rad electrophoresis unit and set it to a constant current of 220mA. Transfer the film for 2 hours. Place the NC membrane in pre-prepared 5% skimmed milk powder containing TBST at room temperature for blocking. Incubate the blocked NC membrane with specific primary and secondary antibodies respectively. Position the NC membrane within the automated chemiluminescence imaging system. Utilise the software to capture chemiluminescent images and retain the raw files. Quantify the results using ImageJ software (NIH, Bethesda, MD, USA; http://rsb.info.nih.gov/nih-image/) to analyse the acquired images.

#### Real-time QPCR

Total RNA was extracted from cardiac tissue or cells using the TransZol™ Up Plus RNAKit (TransGen Biotech, Beijing, China) and reverse transcribed into first-strand cDNA using the RevertAid First Strand cDNA Synthesis Kit (Thermo Fisher Scientific). Real-time PCR amplification was performed using the ABI 7900HT Real-Time PCR System (Thermo Fisher Scientific) and TransStart® Top Green qPCR SuperMix (TransGen Biotech). Primer sequences were synthesised according to Table S2.

#### OGTT

Mice were fasted overnight but maintained normal access to drinking water. Glucose solution was prepared in advance using triple-distilled water as the solvent. A final concentration of 20% glucose solution (w/v) was prepared and filtered to ensure sterility. Tail-prick blood samples were sequentially collected from each mouse, and fasting blood glucose levels were measured using a blood glucose meter, with data recorded. Immediately following blood collection and measurement, each mouse was injected with the pre-prepared glucose solution at a dose of 2 g/kg, with an injection volume of 10 μL per gram of body weight. At 15 minutes, 30 minutes, 60 minutes, 90 minutes, and 120 minutes post-injection, sequential tail-stick blood samples were taken from each mouse. Fasting blood glucose levels were measured using a blood glucose meter, and data were recorded. Maintain a quiet environment during testing to avoid stress-induced blood glucose elevation.

#### Masson’s trichrome staining

Tissues were fixed in 10% formalin and routinely dehydrated and embedded. Mouse hearts were dewaxed prior to staining. Sections were placed in Bouin’s solution overnight at room temperature for enzyme staining, then rinsed under running water until the yellow colouration on the sections disappeared. Sections were sequentially stained with azurine blue for 2-3 minutes, Mayer’s haematoxylin drop-stained for 2-3 minutes, differentiated for several seconds with acidic ethanol, counterstained with Alcian blue solution for 10 minutes, treated with phosphomolybdic acid solution for approximately 10 minutes, stained with aniline blue solution for 5-10 minutes, treated with a weak acid solution for 2 minutes, dehydrated three times with absolute ethanol for 5-10 seconds each, and cleared with xylene three times for 1-2 minutes each. Finally seal the sections with neutral resin.

#### Tissue immunofluorescence

Fix with 4% paraformaldehyde for 6 hours, then dehydrate in 30% sucrose solution for 12 hours until tissue settles. Permeabilise tissue with 0.5% Triton-100 permeabilisation solution for 20 minutes. Block with goat serum blocking solution for 1 hour, then wash three times with PBS for 5 minutes each to remove excess blocking solution. Incubate overnight at 4°C with primary antibody. Incubate with secondary antibody at room temperature, protected from light, for 1 hour. Mount using DAPI-containing fluorescent mounting medium. Wash with PBS between each step.

#### Metabolomics

In brief, add 1 µl of the internal standard mixture (each internal standard containing 5 ng: LTB4-d4, PGE2-d4, 6-keto-PGF1α-d4, 11,12-DHET-d11, 20-HETE-d6, 8,9-EET-d11, 5-HETE-d8, ARA-d8, EPA-d5, DHA-d5, and 9-HODE-d9). Arachidonic acid and free fatty acids were sequentially extracted with methanol (containing 0.01% formic acid) and ethyl acetate under dim light conditions. Samples were evaporated to dryness under a gentle nitrogen stream, redissolved in 30% acetonitrile, and filtered. Extraction tubes employed were bottom-adsorption centrifuge tubes. Chromatographic separation was performed using ultra-performance liquid chromatography (UPLC) on a BEH C18 column (1.7μm, 100 × 2.1mm i.d.). Targeted analysis of arachidonic acid metabolites was performed using a 5500 QTRAP Hybrid Triple Quadrupole-Linear Ion Trap Mass Spectrometer (AB Sciex, Framingham, MA, USA) equipped with a turbo ionisation electrospray ion source. Analysis of raw data was performed using MetaboAnalyst 6.0 (https://www.metaboanalyst.ca/). Missing values were imputed as half the minimum positive value, and all data underwent automatic scaling and log transformation.

#### Primary cardiomyocyte cultures

Within 24 hours of birth, SD rats were dissected via a midline incision at the cardiac region using curved scissors to extract the heart. The heart was minced with scissors, thoroughly rinsed in D-Hank’s solution, and blood clots removed. The minced heart was added to a 10 mL glass vial containing digestion solution (80 mL D-Hank’s solution, pancreatic enzyme 0.085 g, type II collagenase 0.045 g) in a 10 mL glass flask (containing a rotor). Digest and agitate at 37°C for 6 minutes. After settling, transfer the supernatant to a 10 mL centrifuge tube containing DMEM medium supplemented with 10% FBS. Repeat the above steps multiple times until the 10 mL glass vial contains minimal sediment. Pool the liquids from all centrifuge tubes containing supernatant and medium obtained during each step (excluding the first aspirated supernatant). Centrifuge (1000 rpm, 5 minutes, room temperature), collect the cell pellet, filter through a 70μm mesh screen, and collect the filtrate. Transfer the filtered cells to a 10 cm diameter cell culture dish for 1.5 hours. Gently pipette the cells to detach non-adherent cells, which constitute the primary cardiomyocytes. Add 5-bromodeoxyuridine (BrdU) to the primary cell culture medium to inhibit fibroblast formation and purify cardiomyocytes.

#### Omega-3 supplementation

In experiments involving omega-3 supplementation in mice, the omega-3 group was fed a specific grain-based diet supplemented with DHA/EPA, while the control group received a standard grain-based diet without DHA/EPA supplementation (other grain components were identical, with the DHA/EPA proportion compensated for using corn oil). Both diets provided adequate nutrition for normal physiological activity and were nutritionally balanced. The DHA/EPA ratio was determined based on the normal dietary intake range for humans. The specific formulation ratios are detailed in table S3.

#### cAMP assay

Resuspend the sample cells in cell lysis buffer at a concentration of 1 × 10⁷ cells/mL, requiring a minimum of 250μL of cell lysis solution. Determine the expression levels of cAMP within the cells using the cAMP Parameter Assay Kit (R&D Systems, KGE002B).

**Table S1.** Key Resources.

Key Resources Table S1
| REAGENT or RESOURCE | SOURCE | IDENTIFIER |
| --- | --- | --- |
| **Antibodies** |  |  |
| Ki67 Rabbit Monoclonal Antibody | Abcam | Cat#ab16667 |
| PHH3 Rabbit Polyclonal Antibody | Thermo Fisher | Cat#PA5-17869 |
| Aurora B Rabbit Polyclonal Antibody | Abcam | Cat#Ab2254 |
| NR4A1 Rabbit Monoclonal Antibody | Novus Biologicals | Cat#NBP2-66980 |
| PCNA Rabbit Monoclonal Antibody | Cell Signaling Technology | Cat#13110 |
| cTNT Mouse Monoclonal Antibody | Proteintech | Cat#68300-1-Ig |
| Alpha Tubulin Rabbit Polyclonal antibody | Proteintech | Cat#11224-1-AP |
| CREB Rabbit Monoclonal Antibody | Abcam | Cat#ab32515 |
| CREB (phosphorylation) Rabbit Monoclonal Antibody | Abcam | Cat#ab32096 |
| **Critical commercial assays** |  |  |
| cAMP Elisa Kit | R & D | Cat#KGE002B |
| Tranzol up RNA Plus Kit | TransGen Biotech | Cat#ER501-01 |
| Modified Masson's Trichrome Stain Kit | Solarbio Life Science | Cat# G1346 |
| cAMP Elisa Kit | R & D | Cat#KGE002B |
| **Chemicals, peptides, and recombinant proteins** |  |  |
| 17,18EEQ | Cayman Chemical | Cat#50861 |
| 19,20EDP | Cayman Chemical | Cat#10175 |
| 14,15EET | Cayman Chemical | Cat#50651 |
| DAPI | Zsbio | Cat# ZLI-9556 |
| H89 | MCE | Cat#127243-85-0 |
| SQ22536 | MCE | Cat#17318-31-9 |
| 666-15 | MCE | Cat#1433286-70-4 |
| **Bacterial and virus strains** |  |  |
| AAV9-cTnT-GFP | Genechem | YC-A009 |
| AAV9-cTnT-NR4A1 | Genechem | N/A |
| **Experimental models: Cell lines** |  |  |
| H9C2 | Ubigene | N/A |
| **Software and algorithms** |  |  |
| Origin 2019b | Origin | https://www.originlab.com/2019b |
| Prism 9.0 | GraphPad | https://www.graphpad.com/ |
| ImageJ | ImageJ | https://imagej.net/ij/ |
| MetaboAnalyst 6.0 | MetaboAnalyst | https://www.metaboanalyst.ca/ |
| **Oligonucleotides** |  |  |
| Primers used for quantitative RT-PCR | This paper | See Table S2 |

**Table S2.**
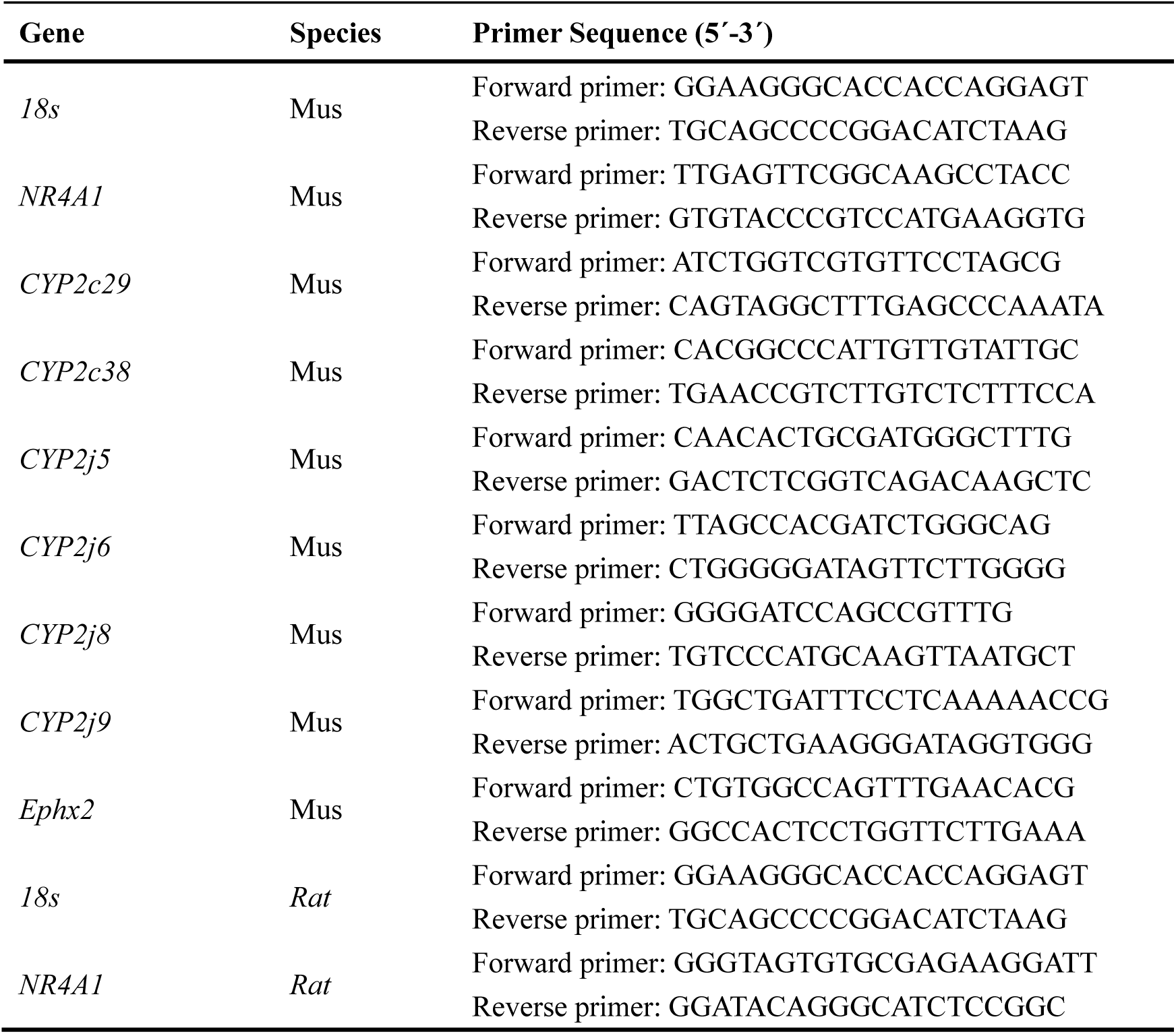
Primer Sequence.

**Table S3.** Omega-3 feed formulation for mice (control group)

| **Ingredients**<br>**(Omega-3)** | **content** | **Ingredients**<br>**(CTRL)** | **content** |
| --- | --- | --- | --- |
| Casein | 18% | Casein | 18% |
| Corn flour | 46% | Corn flour | 46% |
| Sucrose | 23% | Sucrose | 23% |
| Cellulose | 2% | Cellulose | 2% |
| Mineral | 5% | Mineral | 5% |
| Vitamin | 1% | Vitamin | 1% |
| Corn oil | 4.5% | Corn oil | 5% |
| DHA/EPA | 0.5% |  |  |

### Resource Availability

#### Lead Contact

Further information and requests for resources and reagents should be directed to and will be fulfilled by the lead contact, Xu Zhang and Yi Zhu.

#### Materials Availability

All unique/stable reagents generated in this study are available from the lead contact, Xu Zhang, with a completed Materials Transfer Agreement.

---

**Figure S1.**
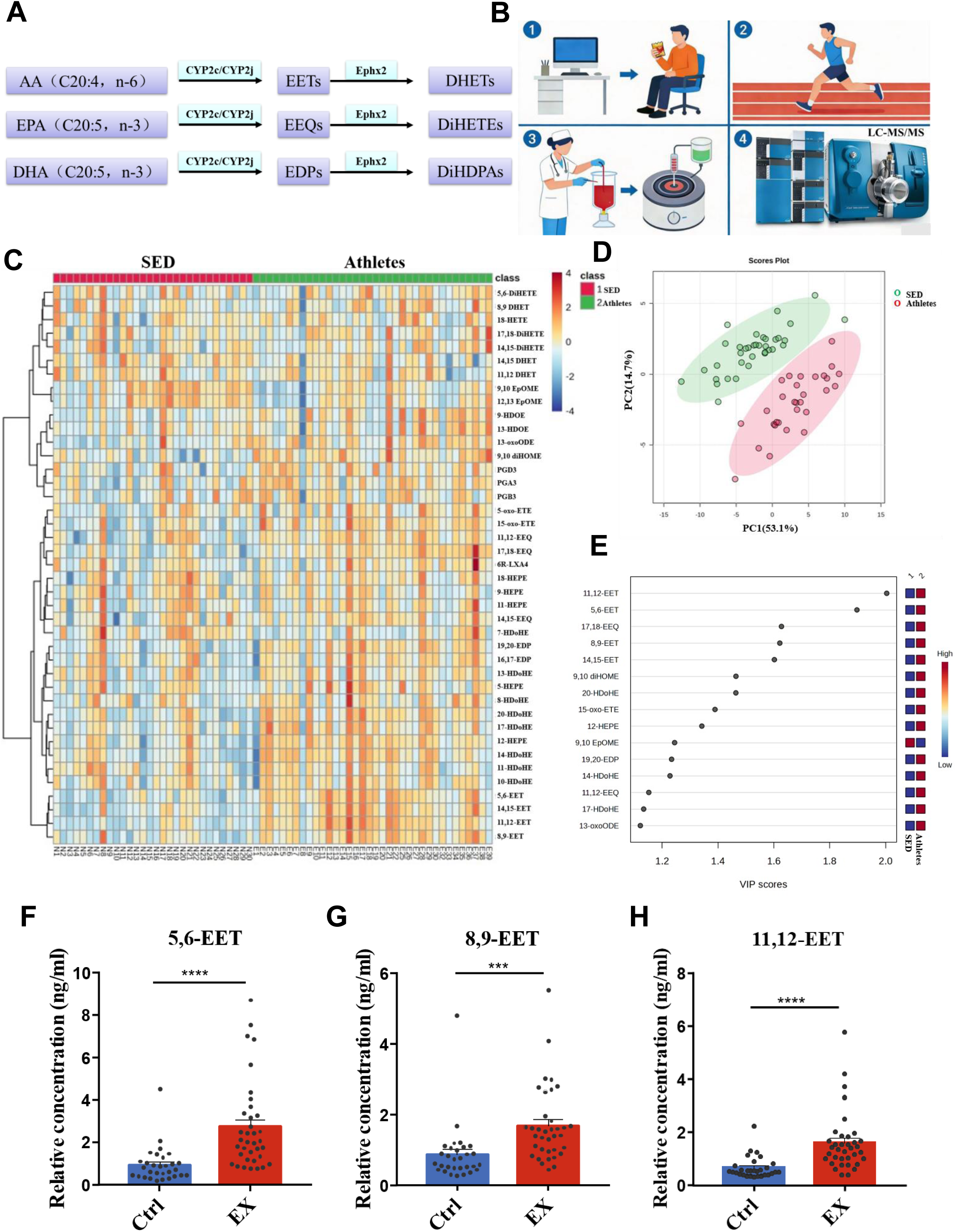
**Eicosanoid profile in elite athletes and sedentary controls.** A. Schematic diagram illustrating the n-6/n-3 PUFA metabolic pathways mediated by CYP450 enzymes. Data are presented as mean ± SEM. *P<0.05; **P<0.01; ***P<0.001. B. Schematic diagram of plasma collection and LC-MS/MS analysis for athletes and sedentary control groups. C. Heatmap visualization of plasma eicosanoid profiles in elite athletes and sedentary controls (age-matched) (Athletes, n=36; Sedentary, n=30). D. Principal component analysis (PCA) score plot showing distinct clustering of metabolite profiles between groups (Athletes, n=36; Sedentary, n=30). E. Variable Importance in Projection (VIP) scores identifying the top 15 metabolites contributing to group separation (Athletes, n=36; Sedentary, n=30). F–H. Plasma concentrations of CYP pathway metabolites 5,6-EET, 8,9-EET and 11,12-EET in elite athletes and sedentary controls. Data are presented as mean ± SEM. *P<0.05; **P<0.01; ***P<0.001; ****P<0.0001.

**Figure S2.**
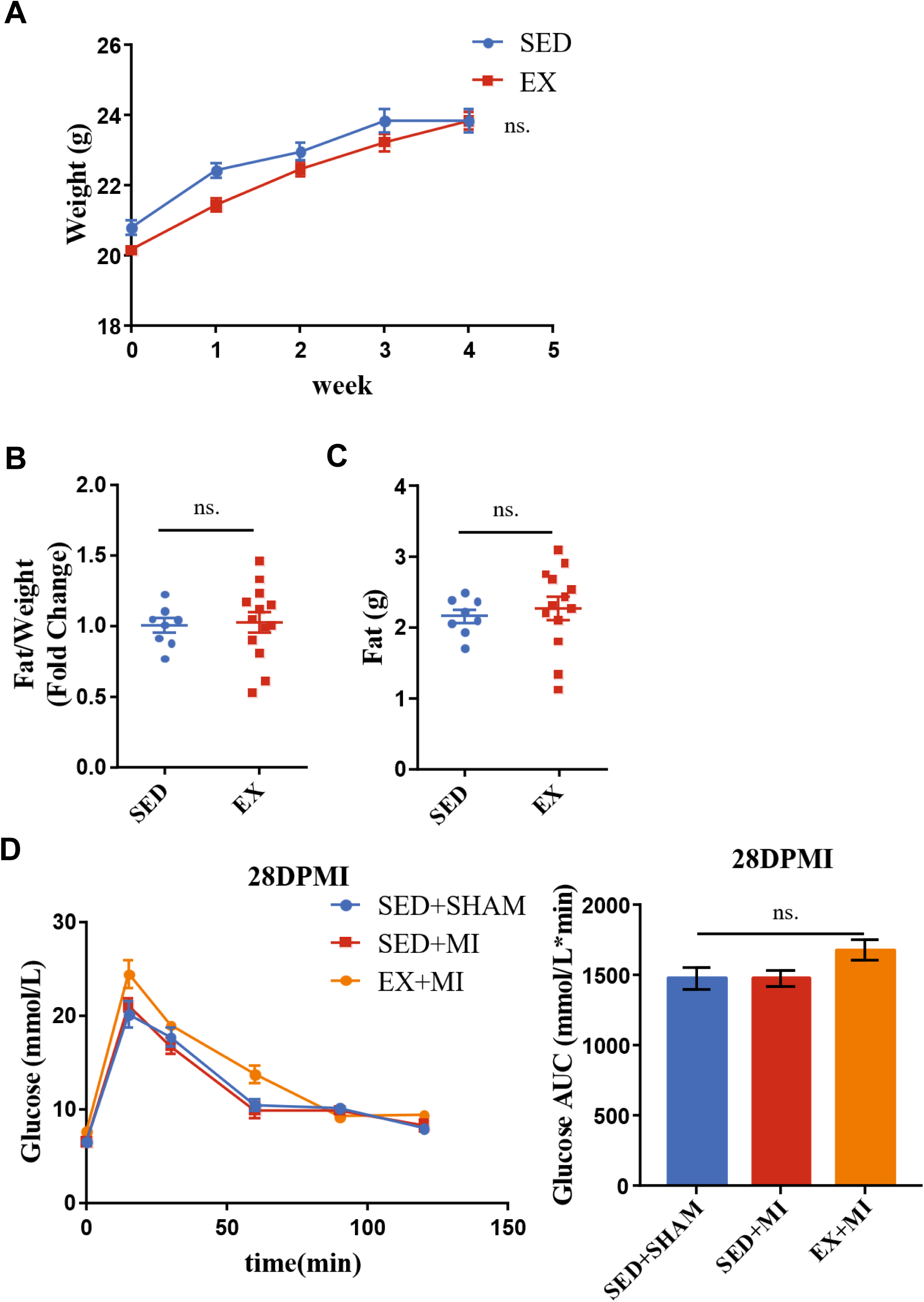
**Impact of aerobic exercise on body composition and glucose metabolism.** A. Body weight of sedentary (SED) and exercised (EX) mice after 28 days of training (SED, n=8; EX, n=13). B-C. Fat-to-body weight ratio (B) and body fat mass (C) assessed by body composition analysis in sedentary and exercised mice (SED, n=8; EX, n=13). D. Glucose tolerance in sedentary sham-operated (SED+Sham), sedentary post-MI (SED+MI), and exercised post-MI (EX+MI) mice (SED+Sham, n=7; SED+MI, n=4; EX+MI, n=6).

**Figure S3.**
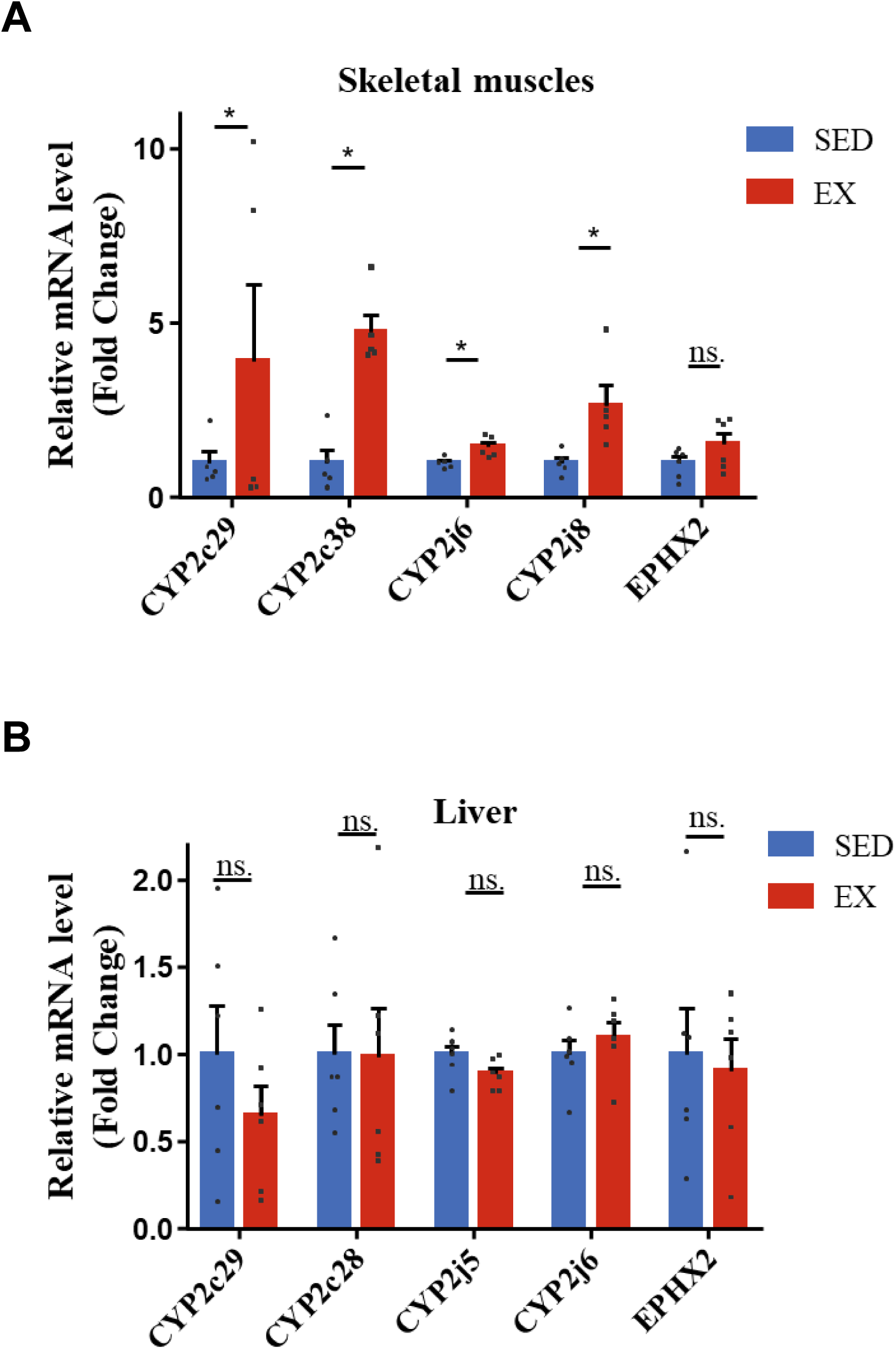
**Effect of aerobic exercise on CYP epoxygenase and EPHX2 expression in peripheral tissues.** (A) Expression levels of cytochrome P450 (CYP) enzymes and EPHX2 in the skeletal muscle of sedentary versus exercised mice (n=5). (B) Expression levels of CYP enzymes and EPHX2 in the liver of sedentary versus exercised mice (n=6). Data are presented as mean ± SEM. *P < 0.05.

**Figure S4.**
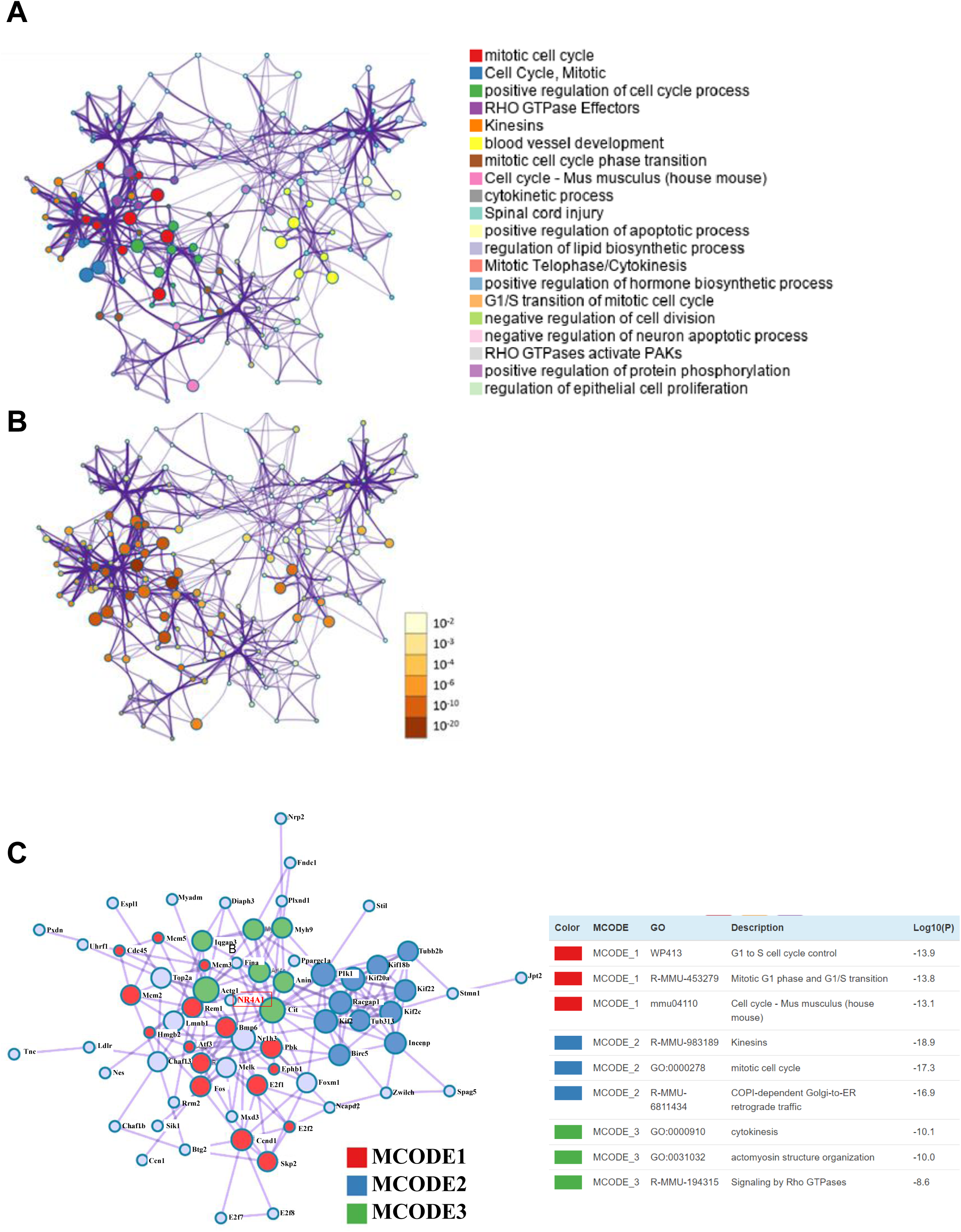
Network analysis of differentially expressed genes in exercised mice post-MI. (A–B) Interaction network of enriched annotations. The diagrams illustrate the correlation between annotations enriched by the differentially expressed genes from the RNA-seq data. The panels on the right display the specific annotation names (A) and their statistical significance levels (B). (C) MCODE analysis of the protein-protein interaction network. The Molecular Complex Detection (MCODE) algorithm was used to identify densely connected topological regions (subnetworks). Functional enrichment analysis reveals that genes within these core subnetworks are primarily enriched in Gene Ontology (GO) terms related to the cell cycle and mitosis.

**Figure S5.**
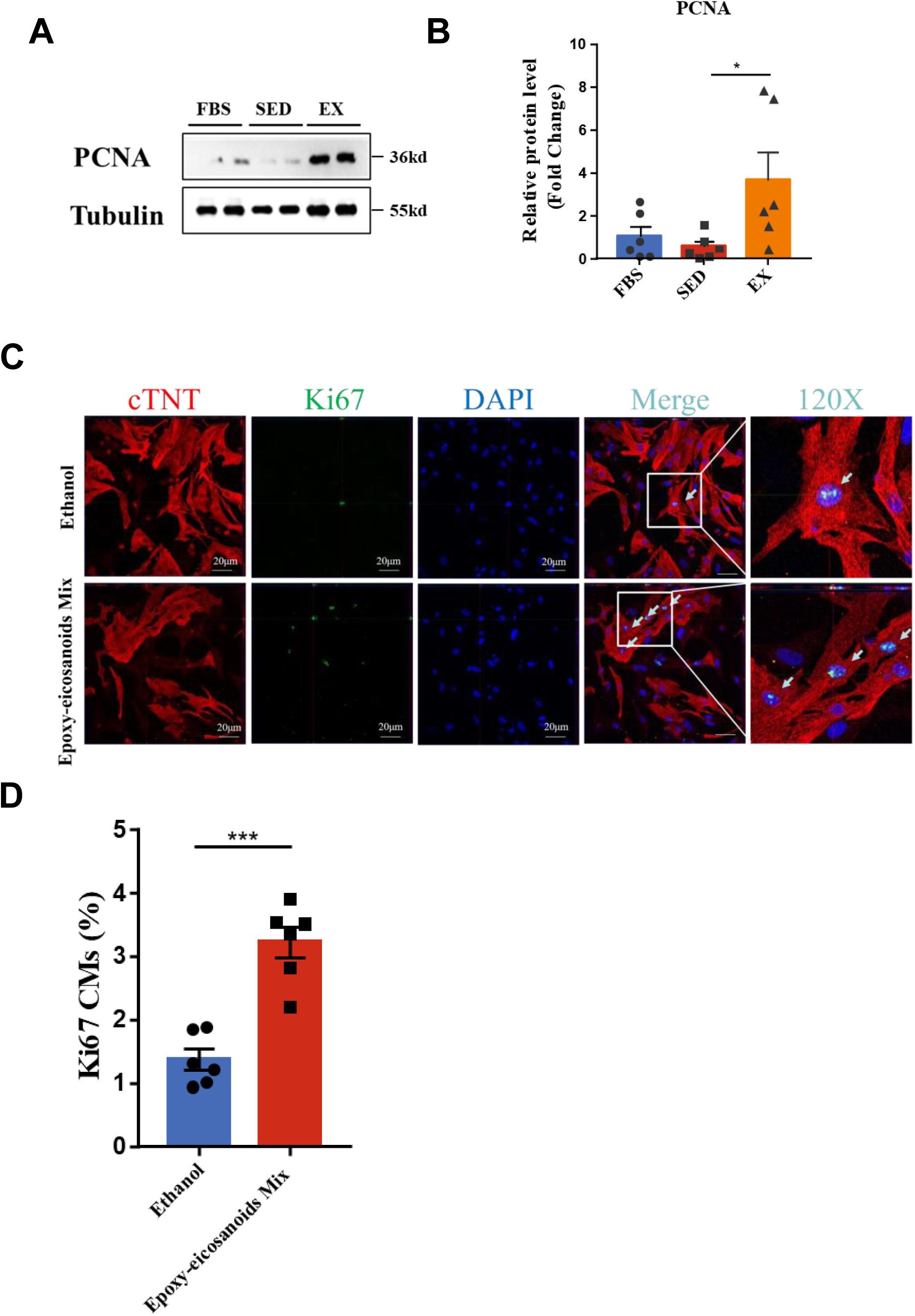
**Exercise-derived lipids and metabolite mixtures promote cardiomyocyte proliferation in vitro.** (A–B) Effect of plasma lipids on proliferation. NRCMs were treated with lipid-enriched plasma fractions from sedentary or exercised mice for 24 hours under hypoxic, serum-free conditions. (A) Representative Western blots and (B) quantification of PCNA protein levels (n=6). (C–D) Effect of a metabolite cocktail on proliferation. NRCMs were treated with a mixture of 17,18-EEQ, 19,20-EDP, and 14,15-EET (or vehicle control) for 24 hours under hypoxic, serum-free conditions. (C) Representative immunofluorescence images and (D) quantification of Ki67-positive cells assessed by High-Content Screening (n=6). Data are presented as mean ± SEM. *P < 0.01, ***P < 0.001.

**Figure S6.**
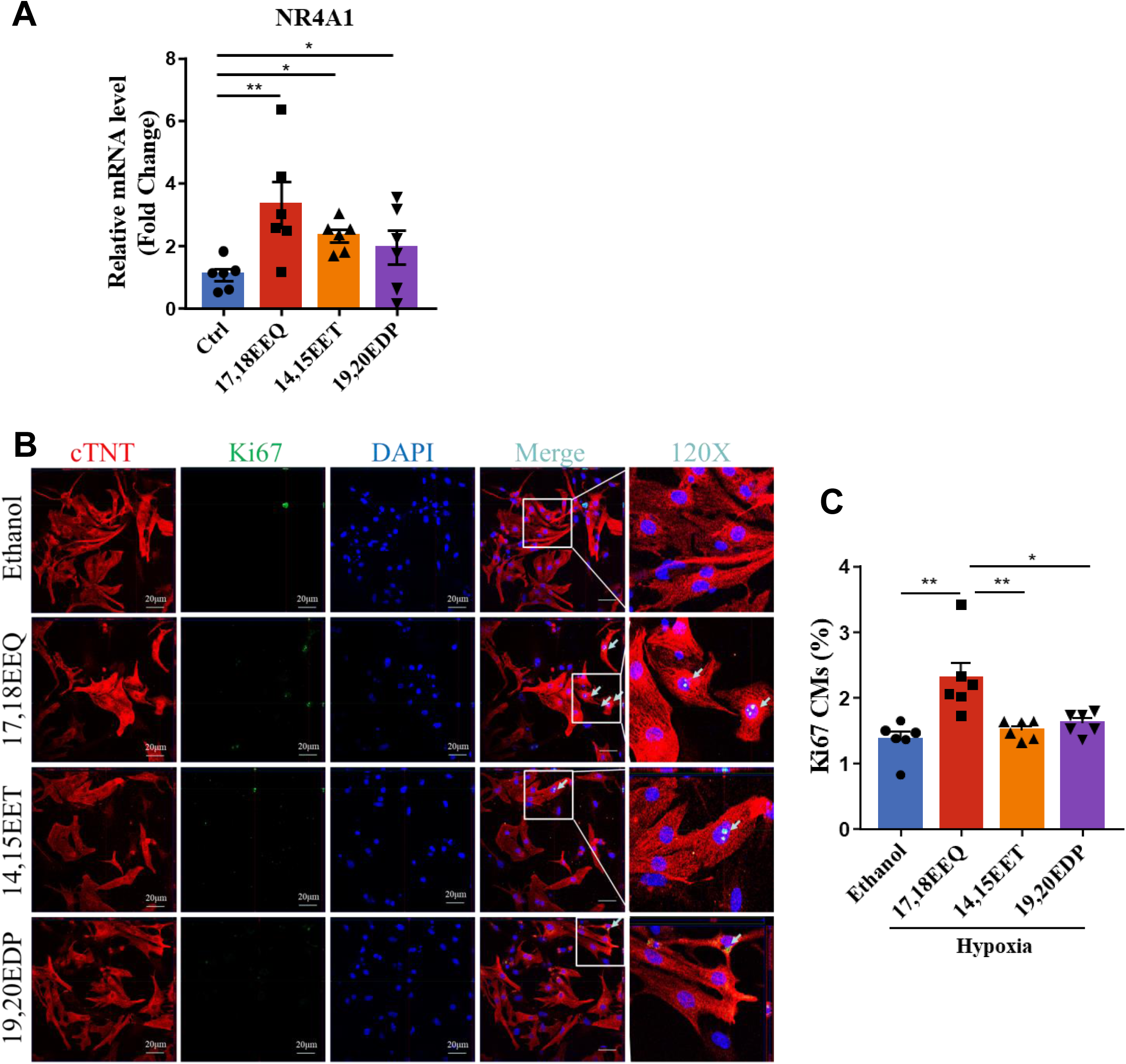
**17,18-EEQ upregulates the exercise-responsive factor NR4A1 and promotes cardiomyocyte proliferation.** (A) mRNA expression levels of Nr4a1 in NRCMs treated with 14,15-EET, 17,18-EEQ, or 19,20-EDP for 24 hours under hypoxic, serum-free conditions (n=6). (B–C) Comparative analysis of cardiomyocyte proliferation. (D) Representative immunofluorescence images of Ki67 and (E) quantification of Ki67-positive cells following 24-hour stimulation with 14,15-EET, 17,18-EEQ, or 19,20-EDP under hypoxic, serum-free conditions. Analysis was performed using High-Content Screening (n=6). Data are presented as mean ± SEM. *P < 0.05, **P < 0.01.

**Figure S7.**
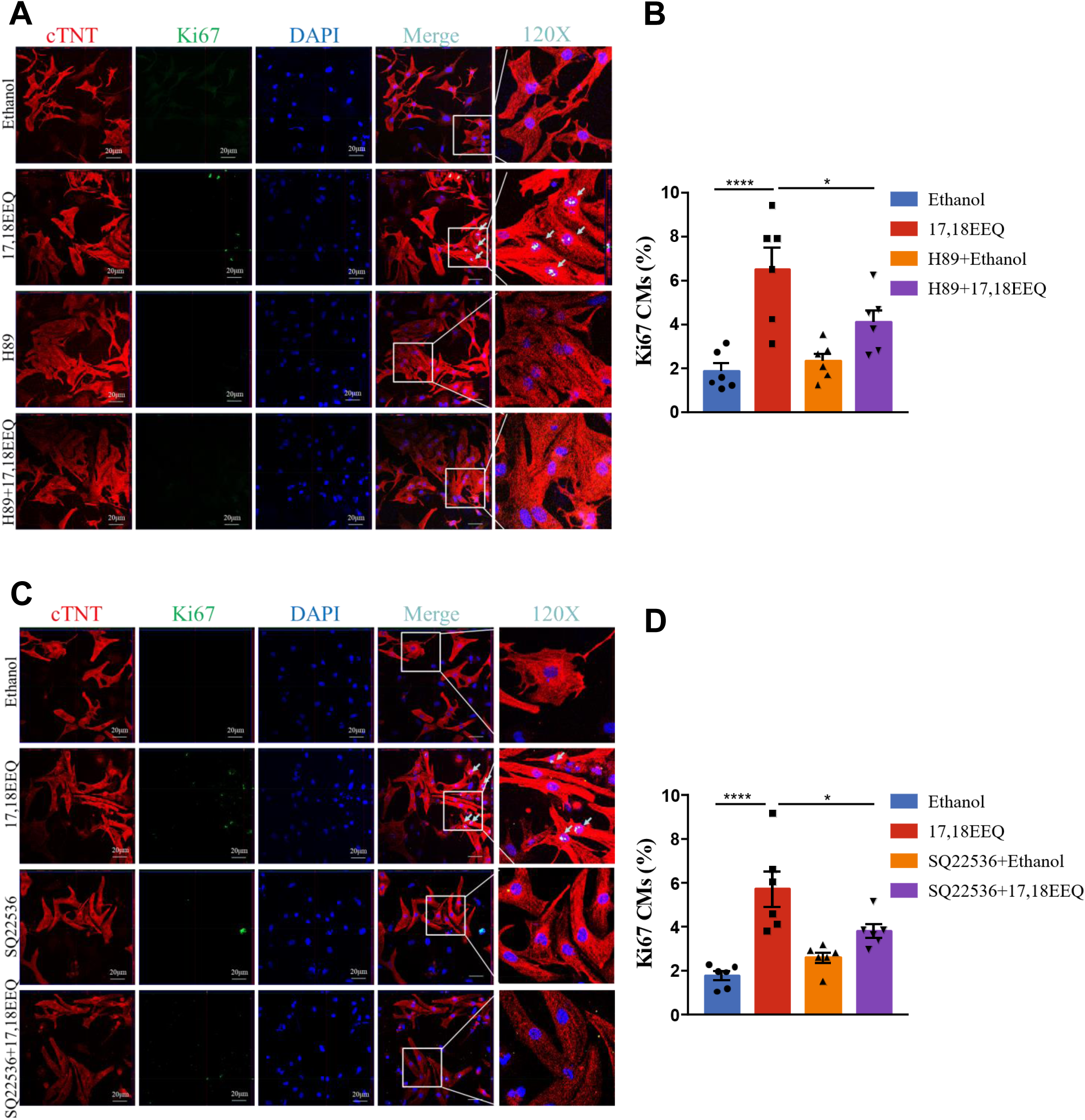
**17,18-EEQ promotes cardiomyocyte proliferation via the cAMP-PKA signaling axis in vitro.** (A–B) Representative immunofluorescence images of Ki67 in NRCMs stimulated with 17,18-EEQ under hypoxic, serum-free conditions. Cells were co-treated with the PKA inhibitor H89 (A) or the adenylyl cyclase inhibitor SQ22536 (B). (C–D) Quantification of Ki67-positive cells assessed by High-Content Screening. Panels show the quantitative analysis corresponding to the H89 (C) and SQ22536 (D) treatment groups (n=6). Data are presented as mean ± SEM. *P < 0.01, ****P < 0.0001.

**Figure S8.**
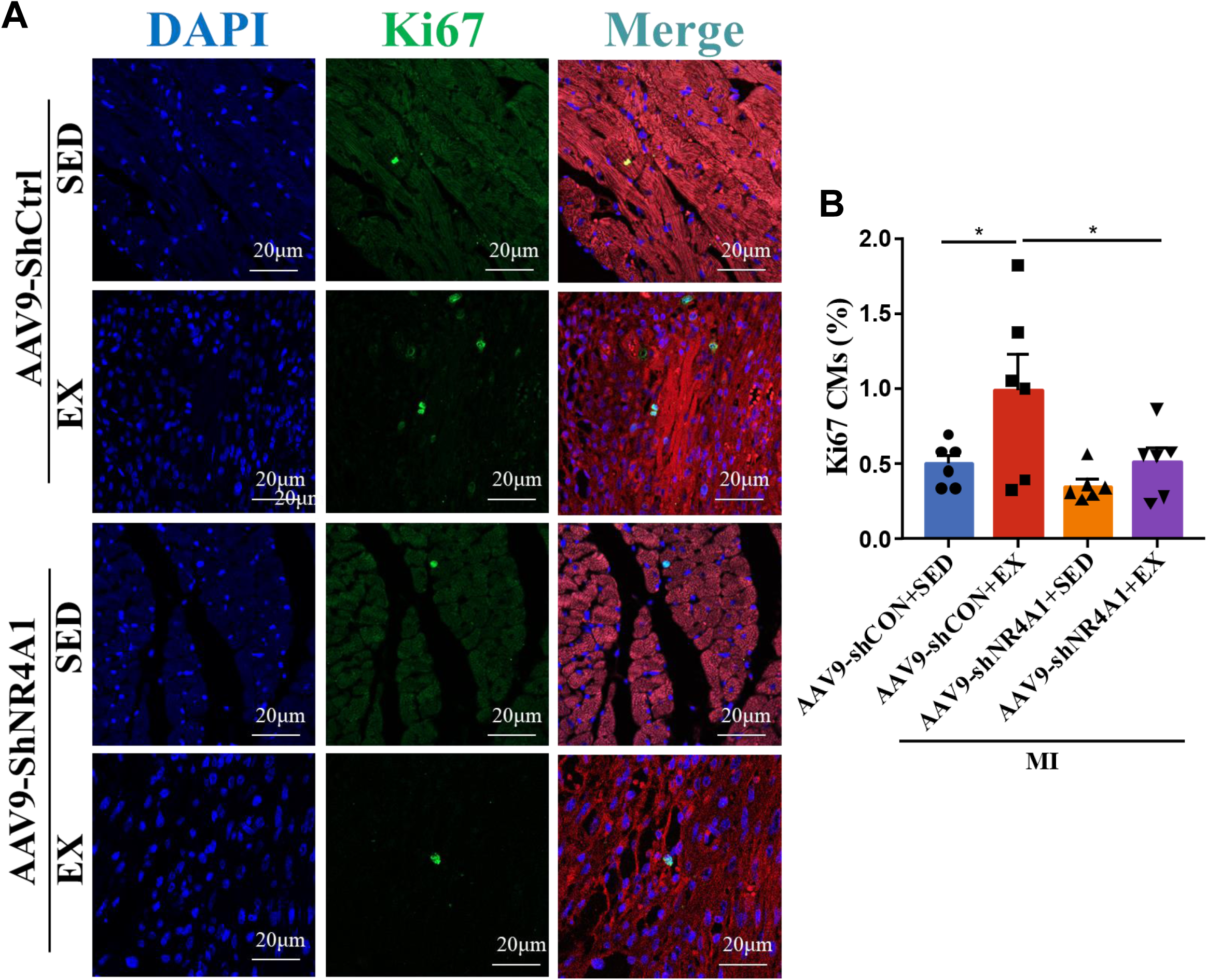
**NR4A1 knockdown abolishes exercise-induced cardiomyocyte proliferation.** (A–B) Evaluation of cardiomyocyte proliferation in vivo. (A) Representative immunofluorescence images of pH3-positive cells in the border zone of the infarct and (B) quantification of pH3-positive cardiomyocytes using High-Content Screening. Mice were subjected to AAV9-mediated NR4A1 knockdown followed by exercise (or sedentary control) and MI (n=6). Data are presented as mean ± SEM. *P < 0.05.

**Figure S9.**
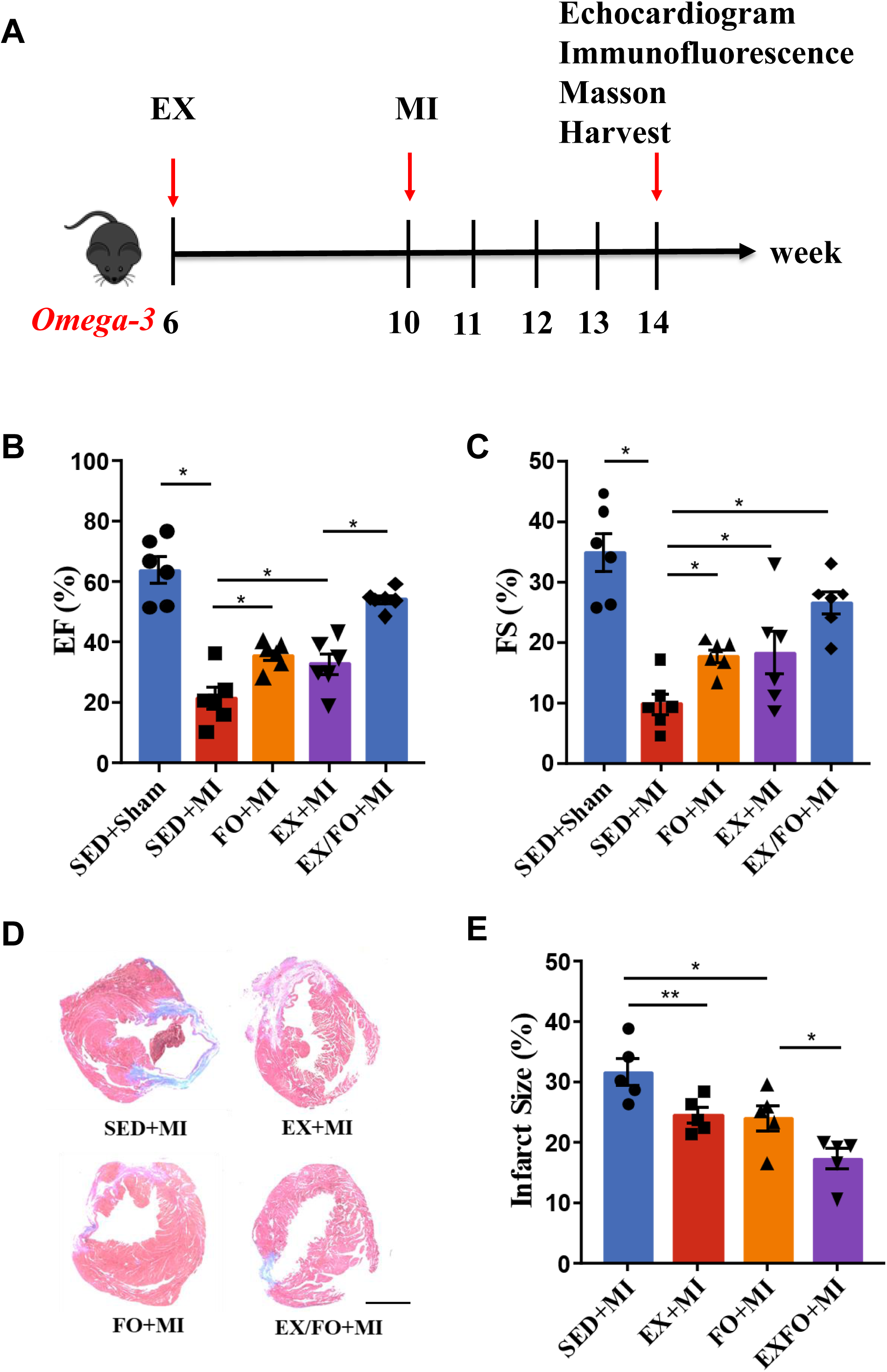
**Aerobic exercise and omega-3 supplementation improve cardiac function and reduce infarct size post-MI.** (A) Schematic illustration of the experimental protocol involving 28 days of aerobic exercise and/or omega-3 supplementation followed by MI induction. (B–C) Echocardiographic assessment of cardiac function. Left ventricular ejection fraction (LVEF) (B) and fractional shortening (LVFS) (C) were measured in mice subject to exercise, omega-3, or combined treatment prior to MI (n=6). (D–E) Assessment of infarct size. (D) Representative Masson’s trichrome staining and (E) quantification of the fibrotic area in the indicated treatment groups (n=5). Data are presented as mean ± SEM. *P < 0.05, **P < 0.01.

**Figure S10.**
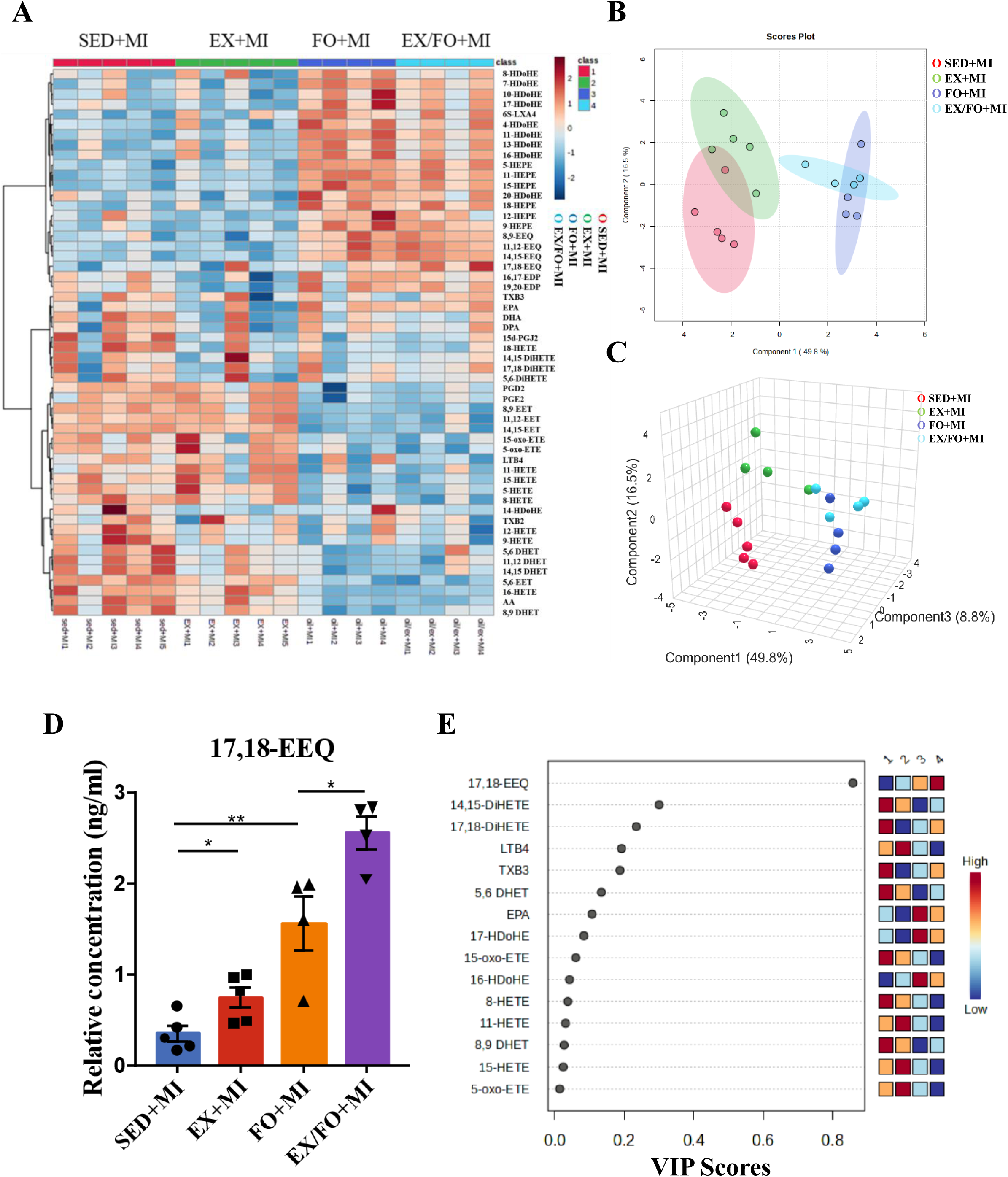
**Effects of aerobic exercise and omega-3 supplementation on the circulating metabolome post-MI.** (A) Heatmap of circulating metabolites in mice undergoing pre-infarction aerobic exercise, omega-3 supplementation, or concurrent aerobic exercise and omega-3 supplementation. (B–C) Sparse partial least squares discriminant analysis (sPLS-DA). (B) Two-dimensional and (C) three-dimensional score plots illustrating the metabolic separation among the groups. (D) Circulating levels of 17,18-EEQ in the indicated groups. (E) Variable Importance in Projection (VIP) score plot identifying the top 15 metabolites contributing to the separation between groups.(SED+MI, n=5; EX+MI, n=5; FO+MI, n=4; EX/FO+MI, n=4). Data are presented as mean ± SEM. *P < 0.05, **P < 0.01.

**Figure S11.**
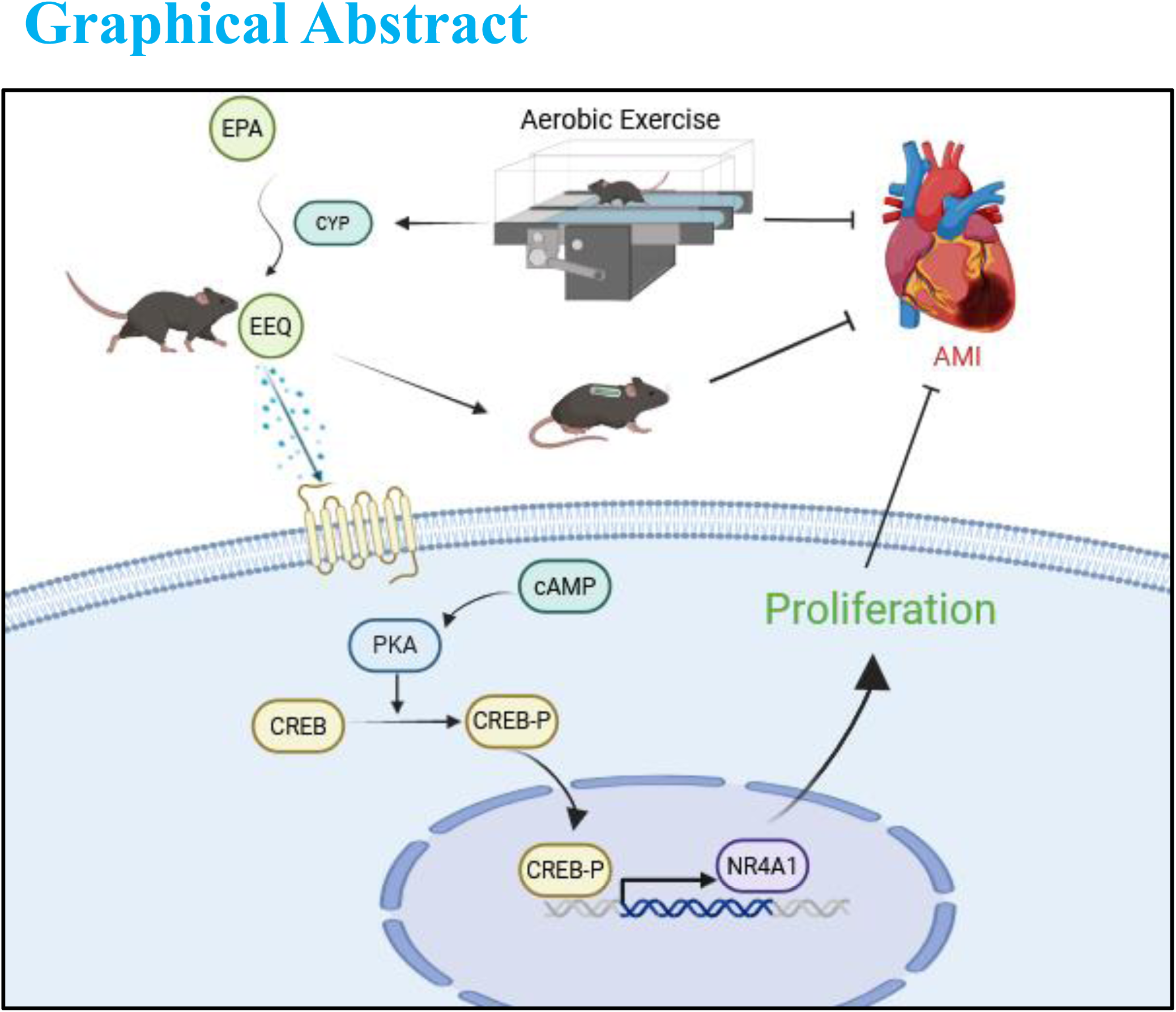

